# Differential roles of cyclin-CDK1 complexes in cell migration and invasion

**DOI:** 10.1101/2024.09.24.614768

**Authors:** Joseph H R Hetmanski, Michael J Jones, Patrick T Caswell, Matthew C Jones

**Affiliations:** Centre for Genome Engineering and Maintenance, Division of Biosciences, Dept of Life Sciences, Brunel University London, London, England; Wellcome Trust Centre for Cell-Matrix Research, School of Biological Sciences, Faculty of Biology Medicine and Health, Manchester Academic Health Science Centre, The University of Manchester, Manchester, United Kingdom.; University of Plymouth, Peninsula Medical School, Plymouth, England

## Abstract

Two key hallmarks of cancer are dysregulated proliferation and metastasis, which are co- ordinated at the single cell level by regulation of cell-cycle progression and invasive cell migration respectively. We have previously described a central role for CDK1 at the nexus of adhesion signalling and cell cycle progression, demonstrating that CDK1 has a non-canonical role in regulating integrin adhesion complexes and in the migration of cancer cells in 3D interstitial matrix. Here we show that the CDK1 binding partners cyclinB1 and cyclinA2 also have roles in cell migration and invasion in both cancer and non-transformed cells. CyclinB1 plays a key role in RhoA activation to promote rear retraction in a membrane tension dependent manner, while cyclinA2 has a general role in promoting motility. Knockdown of either cyclin significantly perturbs migration with contrasting phenotypes, while knockdown of both together has an additive effect which arrests both migration and division. We find that the cell migration specific role of CDK1 is independent of cell-cycle phase, with inhibition or knockdown of CDK1 perturbing migration in G0/G1 arrested cells, while CDK1-cyclin expression correlates strongly with invasive potential of bladder cancer cell lines. Our findings therefore describe how cyclin-CDK1 complexes orchestrate migration as well as division of cells and that cyclinA2-CDK1 and cyclinB1-CDK1 complexes play distinct roles in motility. Furthermore, these findings suggest that targeting CDK1 signalling in aggressive and invasive tumours may have an unexpected dual potential to combat metastasis in addition to proliferation.

## Introduction

Cell migration is a fundamental process that is required for development and physiological processes such as wound healing, angiogenesis and during immune responses(Yamada and Sixt, 2019). Furthermore, invasive cell migration into surrounding tissues is a hallmark of advanced cancer and contributes to metastasis in a variety of settings(Hanahan and Weinberg, 2011; Polacheck et al., 2013). Cell migration requires a concerted regulation of extracellular matrix adhesion, cytoskeletal dynamics and membrane tension that facilitate motility (Seetharaman and Etienne-Manneville, 2020; SenGupta et al., 2021). Enhancing our understanding of the molecular processes that coordinate migration is required to develop an integrated model of how cell migration is regulated in both physiological and pathological situations.

We have recently described a role for CDK1 in regulating integrin adhesion complexes and the actin cytoskeleton during cell cycle progression and migration (Hetmanski et al., 2021; Jones et al., 2018). This regulation occurs in part due to a direct interaction between CDK1 and the adhesion protein talin (Gough et al., 2021), via kinase-dependent regulation of the formin FMNL2 (Jones et al., 2018) and through RhoGTPase-mediated rear retraction (Hetmanski et al., 2021). Several other cytoskeletal targets for CDK1 have been identified, suggesting that regulation of the cytoskeleton is a crucial aspect of the role of CDK1 in cells and may represent the mechanism by which CDK1 controls both motility and division (Jones et al., 2019; Jones and Jones, 2024). In addition to our own studies in ovarian cancer cells migrating in 3D cell-derived matrices, knockdown of CDK1 abrogates cell migration and invasion in breast cancer cells (Wang et al., 2021), cholangiocarcinoma cells (Duan et al., 2021), and hepatocellular carcinoma (Dang et al., 2021), and inhibition of CDK1 activity reduces vimentin phosphorylation and migration in Schwann cells (Chang et al., 2012). This suggests that CDK1 plays a conserved role in regulating cell migration in both normal and cancer cells, however the role for CDK1 in mediating cell migration in 2D and 3D environments remains ill-defined.

CDK1 function is primarily mediated through interaction with partner cyclin proteins that act to guide CDK1-specific phosphorylation (Morgan, 1995). Whilst many functions of CDK1 are associated with the cell cycle, cell cycle independent roles have also been identified including in DNA damage repair (Duda et al., 2016; Esashi et al., 2005; Hentges et al., 2014; Swaffer et al., 2016) and motility (Chang et al., 2012; Dang et al., 2021; Duan et al., 2021; Hetmanski et al., 2021; Wang et al., 2021). CDK1 forms complexes with A- and B-type cyclins (Malumbres, 2014), and regulation of adhesion complexes by CDK1 occurs primarily via association with cyclinA2 (Jones et al., 2018). CyclinA2 expression mediates cell migration, invasion, and metastasis in hepatocellular (Fu et al., 2021), lung (Ruan et al., 2017), and breast carcinomas (Lu et al., 2022). However, low levels of cyclinA2 in prostate (Mashal et al., 1996), colorectal (Guo et al., 2021), and oral squamous cell carcinoma (Wang et al., 2008) is associated with increased invasiveness and more aggressive disease. In normal mouse epithelial cells, loss of cyclinA2 drives epithelial-mesenchymal transition (EMT) and promotes mesenchymal cell migration (Bendris et al., 2014). These findings therefore describe a potential role for cyclinA2 in regulating CDK1 function during cell migration and invasion that is likely to be cell and context dependent. In contrast, little is known with regards to whether cyclinB1 can regulate cell motility, despite cyclinB1-CDK1 complexes being able to modify components of the cell migratory machinery such as focal adhesions, actin, RhoA and microtubule dynamics during mitosis (Jones et al., 2019).

Our previous work has demonstrated that localisation of the RhoA-GEF Ect2 to caveolae at the rear of cells migrating in 3D matrices is dependent on CDK1 activity (Hetmanski et al., 2021). Low membrane tension at the cell rear promotes the formation of caveolae which in turn drive RhoA activity via Ect2 and allow forward movement of the cell rear (Hetmanski et al., 2019). CDK1 is required for cell cycle Ect2 activation, and CDK1 inhibition also abrogates RhoA activity to prevent rear retraction (Hetmanski et al., 2021), indicating that there is significant overlap in the regulation of the cytoskeleton in division and migration. Because both cyclinA2 and cyclinB1 have been shown to regulate RhoA activity (Arsic et al., 2012; Matthews et al., 2012; Niiya et al., 2006), we sought to determine which cyclin-CDK1 complexes were required for the modulation of cell migration in physiologically relevant 3D matrices. We demonstrate that cyclinB1 localises to the rear, whilst cyclinA2 shows a more diffuse localisation throughout the cytoplasm and nucleus of motile cells. Knockdown of either cyclinA2 or cyclinB1, or both simultaneously, perturbs cell motility in cell-derived matrices and invasion in collagen gels in a cell-cycle independent manner in both normal and cancer cells. However, cyclinA2 and cyclinB1 depletion had differing effects on cell morphology, membrane tension, caveolae localisation and GTPase activity suggesting that cyclin-CDK1 complexes have distinct roles in regulating cell migration and invasion with cyclinB1-CDK1 controlling rear retraction and cyclinA2-CDK1 important for protrusion. These findings therefore describe how changes in expression levels of cyclinA2, cyclinB1 and CDK1 or dysregulation of cyclin-CDK1 complexes may impact upon cancer cell invasive migration in addition to proliferation and suggest that targeting CDK1 activity in invasive tumours such as muscle invasive bladder cancer could be explored as a therapeutic option.

## Results

### CyclinA2 and cyclinB1 control different aspects of cell migration in 3D cell-derived matrix

We previously showed that CDK1 is involved in cell migration in 3D environments and functions upstream of RhoA activity and contractile machinery at the rear of the cell (Hetmanski et al., 2021). We therefore investigated the role of the well described CDK1 binding partners cyclinB1 and cyclinA2 in migrating A2780 ovarian cancer cells. Depletion of either cyclinA2 or cyclinB1 by siRNA (Figure S1A) significantly reduced the speed of A2780 ovarian cancer cell migration in 3D cell-derived matrices (CDM), (Figure 1A,B, Movie S1). While cyclinB1-depleted cells were characterised by a longer and narrower shape consistent with a rear retraction defect (Cramer, 2013; Hetmanski et al., 2019) (Figure 1A, Movie S1), the effect of cyclinA2 depletion was strikingly different, with cells displaying a wider and more rounded morphology. We therefore tested rear retraction directly in cells expressing cherry-Cav1 to indicate caveolae localisation to the rear – a key component of the rear retraction positive feedback machinery we previously identified (Hetmanski et al., 2019). While both cyclinA2 and cyclinB1 knockdown decreased rear movement compared to control siRNA (Figure 1C,D), cyclinB1 depletion caused a more pronounced reduction and cells displayed a loss of rear Cav1 localisation, while Cav1 remained localised at the rear of cyclinA2 depleted cells. This suggests that cyclinB1 perturbation has a direct effect on rear contractile machinery, while cyclinA2 knockdown reduces rear retraction perhaps due to an overall decrease in cell speed. CyclinB1 knockdown increased the distance from the retracting rear to the nucleus, whilst cyclinA2 reduced this, (Figure S1B), further indicating that cyclinB1 is required for active translocation of the rear whereas cyclinA2 may prevent influence the position of the nucleus by controlling other aspects of migration such as protrusion. The CDK1-cyclinB effect on rear retraction/migration occurs primarily via cyclinB1, as cyclinB2 siRNA had no significant effect on migration speed while cyclinB1 + cyclinB2 siRNA together had a similar effect on reducing migration speeds as cyclinB1 alone (Figure S1C).

**Figure 1:**
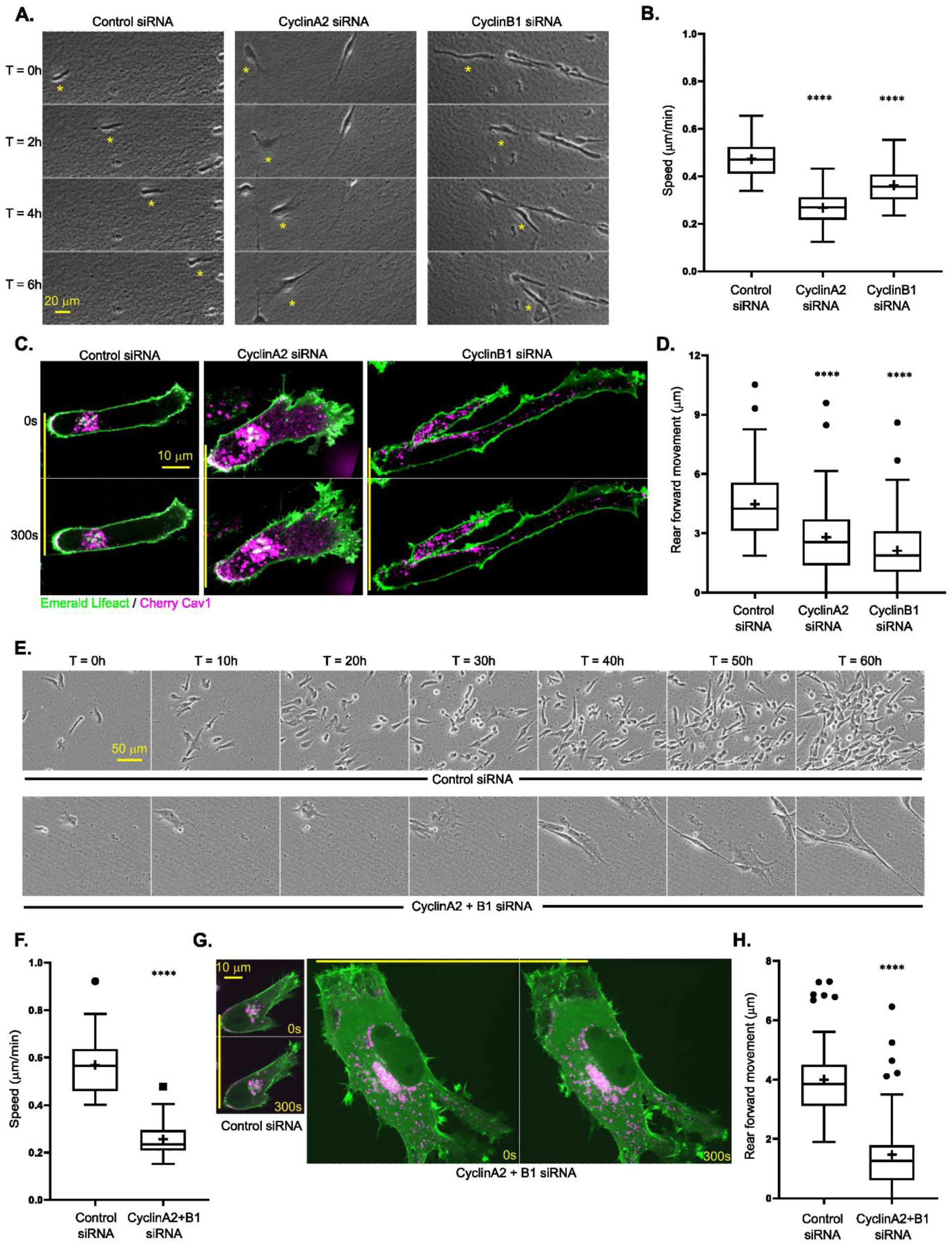
CyclinA2 and cyclinB1 differentially regulate cell migration in 3D cell-derived matrix . **A.** Control, cyclinA2 and cyclinB1 knockdown A2780 ovarian cancer cells seeded in 3D cell derived matrix (CDM) migrating over 6 hours, yellow * denotes the position of the same cell at each time point. **B.** Average migration speed of A2780 cells in CDM across 16 hour timelapse, n = 75 cells per condition analysed across 3 repeats. **C.** Control, cyclinA2 and cyclinB1 knockdown A2780 cells expressing Emerald Lifeact (green) and Cherry Cav1 (magenta) seeded in CDM; solid yellow line indicates initial rear position to compare to position after 300s; cell moving from left to right (rear on the left, leading edge at the front) in this image and throughout. **D.** Average forward rear movement of A2780 cells in CDM across 300s timelapse, n > 40 cells analysed per condition across 3 repeats. **E.** Timelapse images of Control and cyclinA2 + cyclinB1 concomitant knockdown A2780 cells seeded in CDM over 60 hours, same position for each frame and same scale for each condition. **F.** Average migration speed of A2780 cells in CDM across the first 16 hours of timelapse, n = 75 cells per condition analysed across 3 repeats. **G.** Control and cyclinA2 + cyclinB1 concomitant knockdown A2780 cells expressing Emerald Lifeact (green) and Cherry Cav1 (magenta) seeded in CDM, yellow line denotes initial rear position for comparison purposes, same scale for both images. **H.** Average forward rear movement of A2780 cells in CDM across 300s timelapse, n > 40 cells analysed per condition across 3 repeats. One way ANOVA compared to control used in B and D, unpaired student t-test used in F and H, **** denotes p < 0.0001 throughout.

Given the significant but phenotypically opposite effects of cyclinA2 and cyclinB1 perturbation on cell motility, we next depleted both cyclins together (Figure S1A). Double knockdown of cyclinA2 + B1 in cells caused a striking phenotype, whereby cells were not only unable to divide over 16 or even 60 hour timelapse acquisitions (Figure 1E, Movie S2), but were also almost completely unable to migrate during those time frames whilst continuing to increase in size (Figure 1E,F, Movie S2). CyclinA2 + B1 knockdown cells displayed hallmarks of the phenotypes seen when cyclinA2 or cyclin B1 were depleted individually: caveolae at the rear were lost and rear forward movement suppressed, similar to cyclinB1 knockdown, while cells also showed wider protrusions, reminiscent of cyclinA2 knockdown (Figure 1G,H). Taken together, these data show that both cyclinA2 and cyclinB1 perturbation have severe effects on cell migration in different ways, while concomitant perturbation of both leads to a striking loss of migratory as well as division capacity.

### CyclinA2 and cyclinB1 differentially regulate cell morphology and RhoA activity in cells within 3D collagen hydrogels

Building on the differing phenotypic effects of cyclin knockdowns on A2780s in CDMs, which are a confined 3D environment (∼20μm thick such that cells are unable to move up and down in Z), we next tested if such diverse responses were present a fully 3D-matrix environment. We seeded control or cyclin siRNA iRFP670 Lifeact expressing A2780s directly into soft collagen gels before analysing cell morphology after 16 hours (Figure 2A). Similar to the CDM environment, cyclinB1 depleted cells appeared long and narrow, with a significantly increased aspect ratio (length / width) in comparison to control cells, while cyclinA2 siRNA cells were significantly wider, bigger overall (as defined by length x width) and had a larger nucleus. Again, an additive effect of both cyclinA2+B1 depletion was observed, whereby cells were considerable larger, longer and wider with larger nuclei than in control, cyclinA2- or cyclinB1-depleted cells (Figure 2B).

**Figure 2:**
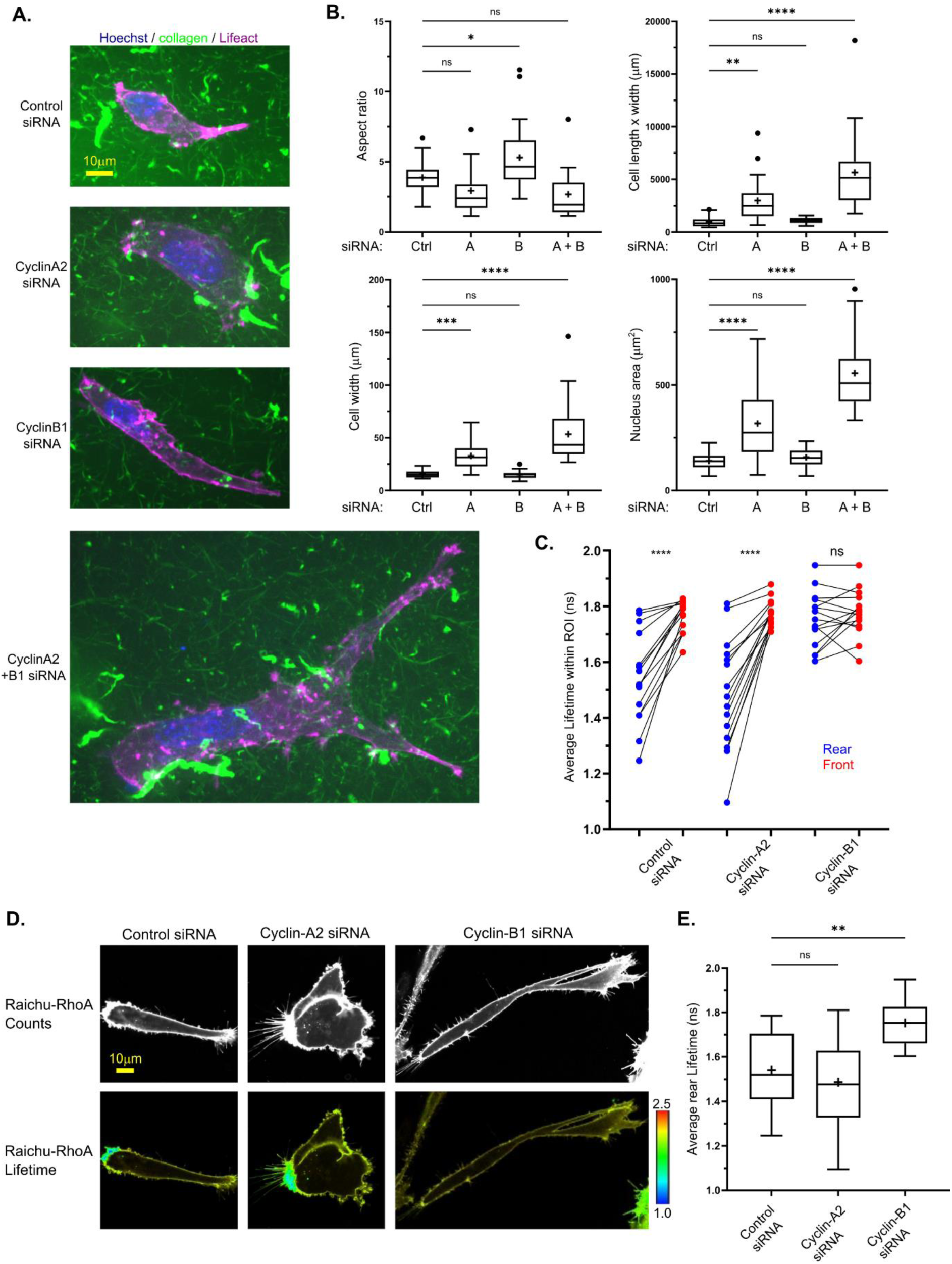
CyclinA2 and cyclinB1 depletion have distinct effects on cell and RhoA activity. **A.** Control, cyclinA2, cyclinB1, and cyclinA2 + cyclinB1 concomitant knockdown A2780 cells stably expressing iRFP670 Lifeact and dyed with Hoechst seeded directly into 3D soft (∼1.5mg/ml) collagen, where 1/10 collagen fibers are labelled with GFP, same scale used for all conditions. **B.** Quantification of cell size and shape metrics of Control (Ctrl), CyclinA2 (A), CyclinB1 (B) and CyclinA2 + CyclinB1 (A+B) siRNA cells; All measurements of length and width taken at widest and longest positions, Aspect ratio (top left) calculated as maximum cell length divided by maximum cell width, n > 25 cells analysed per condition across 3 repeats, same cells analysed for all 4 metrics/graphs. **C.** Average FLIM lifetime of Raichu-RhoA expressing A2780 cells in manually drawn rear (blue) and front (red) regions of interest (rear/front regions of the same cell joined by black line), n > 15 cells analysed per condition across 3 repeats. Note the lower lifetime denotes higher RhoA activity. **D.** Control, cyclinA2, and cyclinB1 knockdown A2780 cells seeded in CDM transfected with GFP-RFP Raichu-RhoA probe, counts (top) and lifetime (bottom) of donor GFP channel shown, colour code lifetime range shown, values in ns: blue denotes shorter lifetime (high activity), yellow/red denotes higher lifetime (low activity). **E.** Average lifetime in rear ROI (same A2780 cells as in C analysed), n > 15 cells analysed per condition across 3 repeats. One way ANOVA compared to control used in all graphs in B and E, paired student t-tests used in C, **** denotes p < 0.0001, *** denotes p < 0.001, ** denotes p < 0.01, ns denotes p > 0.05 (not significant) throughout.

We next explored whether the different morphologies and phenotypes observed corresponded with different RhoA GTPase activity. In CDM we found that cyclinA2 knockdown had little effect on RhoA polarisation or activity at the rear, while cyclinB1 knockdown significantly decreased RhoA activity as measured by an increase in the fluorescence lifetime of the Raichu-RhoA probe specifically at the rear leading to an overall loss of front-rear RhoA polarity (Figure 2C-E). CyclinA2+B1 siRNA together also abrogated front-rear RhoA polarity via a decrease of RhoA activity at the rear (Figure S2A). This further supports the hypothesis that cyclinB1 has a specific role upstream of RhoA and contractile machinery at the rear, while cyclinA2 affects cell morphology and migration via an alternative mechanism, potentially through the regulation of integrin adhesion complexes and talin function in cell protrusions.

### CyclinB1 both responds to and effects membrane tension at the rear

We have found previously that membrane tension plays a key role in driving rear retraction as part of a positive feedback loop (Hetmanski et al., 2019). We therefore tested how cyclin perturbation affected membrane tension using FLIM of the FlipperTR (Colom et al., 2018; Pandzic et al., 2022) in 3D collagen hydrogels. As we found previously in CDM (Hetmanski et al., 2019), A2780 cells displayed lower FlipperTR lifetime at the rear than at the front, indicating lower tension at the rear (Figure 3A, B). Perturbation of either cyclinA2 or cyclinB1 individually or both together all resulted in a loss of this membrane tension differential (Figure 3A, B), however only cyclinB1 siRNA specifically increased membrane tension at the rear compared to control cells (Figure 3C). This suggests that cyclinB1 exerts a more direct effect on membrane tension at the rear, while cyclinA2 or cyclinA2+B1 affect membrane tension polarity by altering tension of the membrane more globally via changes in cell shape. or indirectly via the overall decrease in motility which breaks the membrane tension – contractility positive feedback loop.

**Figure 3:**
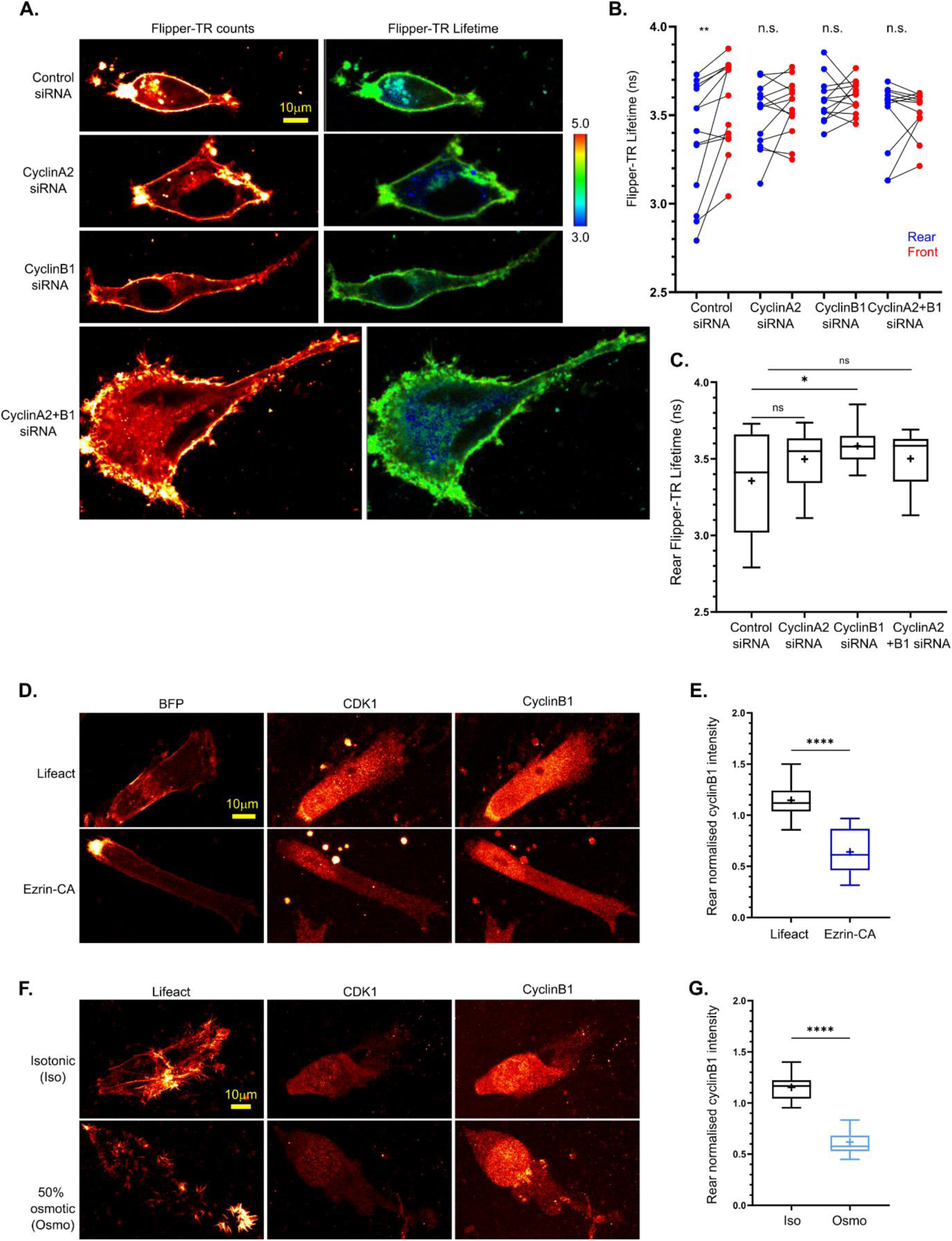
CyclinB1 maintains low membrane tension at the cell rear in migrating cells. **A.** Control, cyclinA2, cyclinB1, and cyclinA2 + cyclinB1 concomitant knockdown A2780 cells seeded directly into soft collagen and dyed with membrane tension probe FlipperTR, counts (left, red hot look-up table LUT applied) and lifetime (colour coded lifetime range shown, values in ns: blue/green denote shorter lifetime – lower tension, yellow/red denote longer lifetime – higher tension) of 488 excitation, 575-625 emission shown. **B.** Average Flipper-TR lifetime of manually drawn rear (blue) and front (red) regions of interest in A2780 cells in collagen gels (rear/front regions of the same cell joined by black line), n > 10 cells analysed per condition across 3 repeats. **C.** Average Flipper-TR lifetime in rear ROI of Control , cyclinA2 (A), cyclinB1 and cyclinA2 + cyclinB1 siRNA cells (same A2780 cells as in B analysed), n > 10 cells analysed per condition across 3 repeats. **D.** Fixed A2780 cells seeded in CDM transfected with BFP Lifeact (top) or BFP Ezrin-CA (constitutively active, to increase membrane tension) and immuno-stained with CDK1 and cyclinB1, red hot LUT applied. **E.** Normalised average cyclinB2 staining intensity at the rear in cells transfected with BFP Lifeact or Ezrin-CA, n > 15 cells analysed per condition across 3 repeats. **F.** Fixed A2780 cells seeded in CDM in normal isotonic medium (top) or subjected to osmotic shock by addition of 50% water for 30 minutes prior to fixation (bottom, to increase membrane tension) and immuno-stained with CDK1 and cyclinB1, red hot LUT applied. **G.** Normalised average cyclinB2 staining intensity at the rear in cells in normal (Iso) or 50% water (Osmo) media, n > 15 cells analysed per condition across 3 repeats. Paired student t-tests used in B, unpaired student t-tests used in E and G, one way ANOVA compared to control used in C, **** denotes p < 0.0001, ** denotes p < 0.01, * denotes p < 0.05, n.s. denotes p > 0.05 (not significant).

Our data so far indicate a specific role for CDK1-cyclinB1 in regulating membrane tension and RhoA activity at the retracting rear (Figures 1,2, (Hetmanski et al., 2021)), therefore we next wanted to ascertain the endogenous localisation of CDK1 and cyclinB1, and whether this was membrane tension dependent. Migrating A2780s in CDM displayed localisation of both CDK1 and cyclinB1 at the rear (Figure 3D, F, Figure S3A, B) when cells were unperturbed or expressing BFP lifeact in normal isotonic medium. When membrane tension was globally increased either by BFP-Ezrin constitutively active (CA) expression or osmotic shock with 50% water, the rear localisation of cyclinB1 and CDK1 was lost (Figure 3D-G) and instead they were both confined more to the nuclear compartment. CDK1 localisation was also more confined to the nucleus in either cyclinB1 or A2 or both together siRNA cells (Figure S3A,B). These data indicate that rear membrane CDK1 localisation is dependent on the availability of cyclin binding partners and that in conditions where this is perturbed CDK1 remains in the nucleus. Moreover, these data suggest that CDK1-cyclinB1 signalling at the rear is both membrane tension dependent and directly affects membrane tension at the rear, indicating CDK1-cyclinB1 is involved in the rear retraction positive feedback loop.

### CyclinA2 and B1 are required for migration of non-cancer RPE cells

Having demonstrated that cyclinA2 and cyclinB1 regulate the motility of cancer cells, we next wanted to test whether these cyclins have conserved roles in regulation of cell migration, morphology, RhoA activity and membrane tension in a non-cancer cell line. Immortalised Retinal Pigment Epithelial (RPE) cells are non-transformed and near diploid, move and divide freely, and have been used extensively in cell cycle studies (Barbiero et al., 2022; Trotter and Hagan, 2020). In RPE cells, knockdown of cyclinA2, cyclinB1 or both cyclinA2+B1 (Figure S4A) all severely reduced migration in 3D CDM (Figure 4A,B), where cyclinB1 depleted cells were longer and thinner cells, cyclinA2 depleted cells were wider and larger overall, and cyclinA2+B1 knockdown together showed an additive effect where cells were largest and moved the least. Similar effects were seen in 3D collagen gels (Figure 4C) where cyclinA2 or cyclinA2+B1 knockdown cells also displayed many more leading edge protrusions, while cyclinB1 knockdown resulted in a loss of rear RhoA activity and cyclinA2+B1 siRNA led to a mis-localisation of RhoA activity (Figure 4C). For RPE cells in 3D collagen, knockdown of either cyclin individually or both together resulted in the loss of front-rear membrane tension polarity while only cyclinB1 siRNA specifically increased membrane tension at the rear (Figure 4D-F). Altogether these data illustrate that the phenotypic consequences of cyclinA2 and cyclinB1 siRNA observed previously in ovarian cancer A2780s are conserved in non-cancer RPEs, whereby during motility cyclinB1 is involved in rear retraction while cyclinA2 is involved elsewhere.

**Figure 4:**
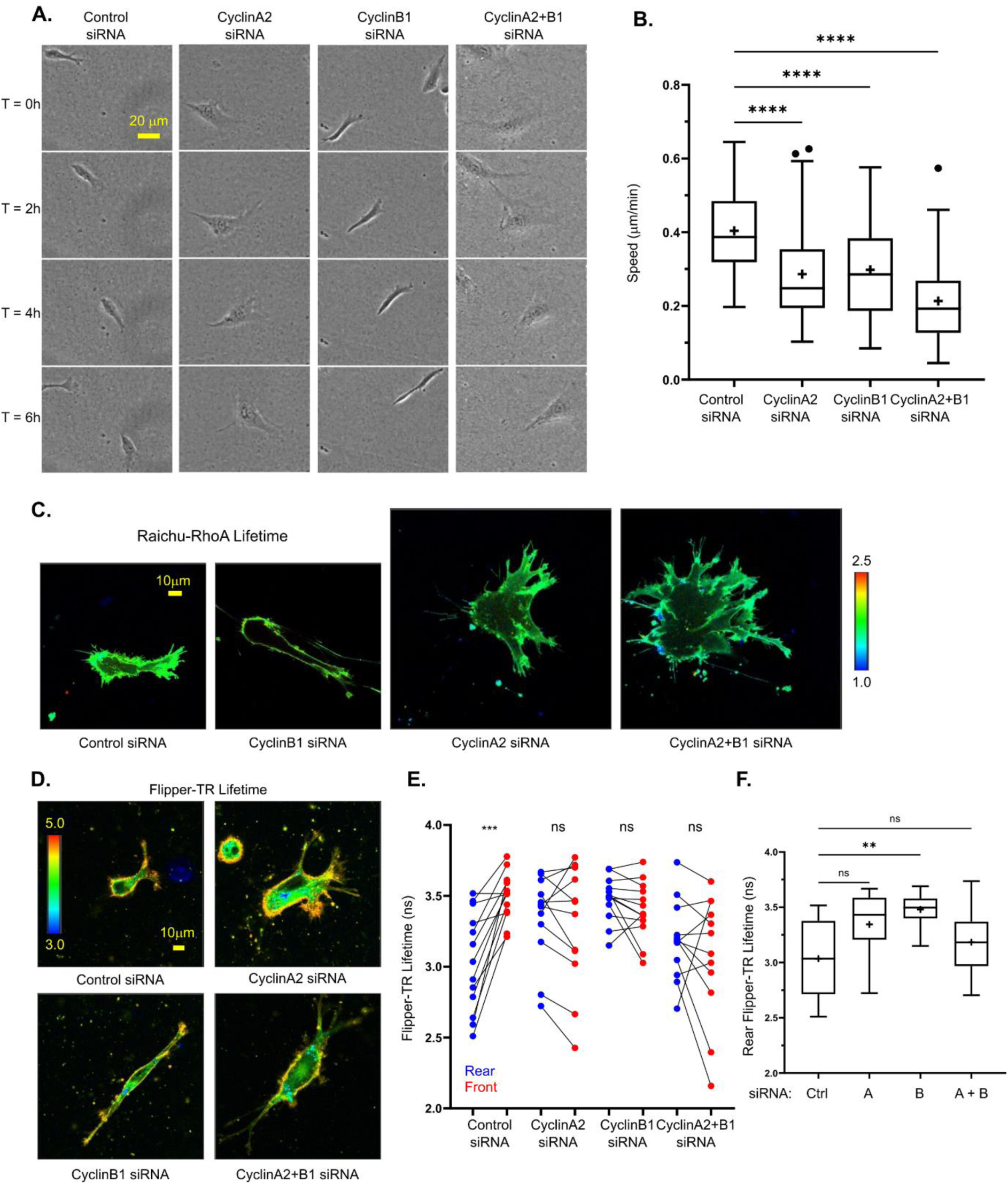
Cyclin perturbation in non-cancer RPE cells demonstrates conserved differential roles for cyclinA2/B1 in invasive migration. **A.** Control, cyclinA2, cyclinB1, and cyclinA2 + cyclinB1 concomitant knockdown retinal pigment epithelial (RPE) cells seeded in CDM migrating over 6 hours, the same cell moving from left to right in each frame shown. **B.** Average migration speed of RPE cells in CDM across 16 hour timelapse, n = 75 cells per condition analysed across 3 repeats. **C.** Control, cyclinA2, cyclinB1, and cyclinA2 + cyclinB1 concomitant knockdown RPE cells seeded in CDM transfected with GFP-RFP Raichu- RhoA probe, lifetime of donor GFP channel, colour code lifetime range shown, values in ns: blue denotes shorter lifetime (high activity), green/yellow denotes higher lifetime (low activity). **D.** Control, cyclinA2, cyclinB1, and cyclinA2 + cyclinB1 concomitant knockdown RPE cells seeded directly into soft collagen and dyed with membrane tension probe FlipperTR, lifetime of 488 excitation, 575-625 emission (colour code lifetime range shown, values in ns: blue/green denote shorter lifetime – lower tension, yellow/red denote longer lifetime – higher tension). **E.** Average Flipper-TR lifetime of manually drawn rear (blue) and front (red) regions of interest in RPE cells in collagen gels (rear/front regions of the same cell joined by black line), n > 10 cells analysed per condition across 3 repeats. **F.** Average Flipper-TR lifetime in rear ROI of Control (Ctrl), CyclinA2 (A), CyclinB1 (B) and CyclinA2 + CyclinB1 (A+B) siRNA cells (same RPE cells as in B analysed), n > 10 cells analysed per condition across 3 repeats. Paired student t-tests used in E, one way ANOVA compared to control used in B and F, **** denotes p < 0.0001, *** denotes p < 0.001, ** denotes p < 0.01, n.s. denotes p > 0.05 (not significant).

### Cyclin-CDK1 complexes regulate invasion in ovarian and bladder cancer cells

Given that CDK1-cyclin complexes control the invasive migration of normal and cancer cells, we next tested their role in long range invasion through 3D collagen-I gels. Knockdown of CDK1 or cyclinA2+B1 together resulted in significantly lower invasion over 72 hours in either soft (∼1.5mg/ml) or stiff (∼5mg/ml) collagen (Figure S5A). We next sought to corroborate the role for cyclin-CDK1 complexes in mediating invasive migration in an alternative cancer type. Patients with bladder cancer (BC) tumours that have invaded the surrounding muscle have poor survival rates in comparison to non-muscle invasive tumours and limited treatment options, with the mechanism that contribute to invasive migration by BC cells remining ill-defined (Smith et al., 2024). Initially, we compared expression levels of CDK1, cyclinA2 and cyclinB1 between primary bladder epithelial cells (HUCs), non-invasive BC cells (RT4; Figure S5B) and invasive BC cells (J82, T24 and UMUC3). Expression of CDK1, cyclinA2 and cyclinB1 were all upregulated in the three invasive cell types compared to the two non-invasive lines by western blot (Figure 5A), however this differential expression was not a consequence of drastically increased cell proliferation as all cancer cells proliferated at similar rates (Figure5B and C). We took forward the T24 cells for further analysis as they show enhanced levels of invasion (Figure S5B), and demonstrated that they too show rear localised caveolae, rapid rear forward movement (Figure S5C) and high RhoA activity localised specifically to the rear (Figure S5D). In spheroid invasion assays, T24 cells displayed a dose dependent reduction in invasive capacity through 2.5mg/ml collagen-I gels in response to CDK1 inhibition with RO-3306 up to 5μM (Figure 5D). T24 cell invasion was also reduced in a dose-dependent manner following siRNA-mediated knockdown of CDK1 (Figure 5E) and upon knockdown of either cyclinA2 or cyclinB1 individually, with cyclinA2 siRNA having a stronger reductive effect despite T24 cells demonstrating a compensatory increase in cyclinB1 levels following cyclinA2 knockdown (Figure S5F), while cyclinA2+B1 siRNA together had an additive effect (Figure 5F). These data show that cyclin-CDK1 complexes play an important role in facilitating invasive migration in both ovarian and bladder cancer cells and that changes in expression of CDK1 and partner cyclins may contribute to the acquirement of an invasive phenotype in BC cells.

**Figure 5:**
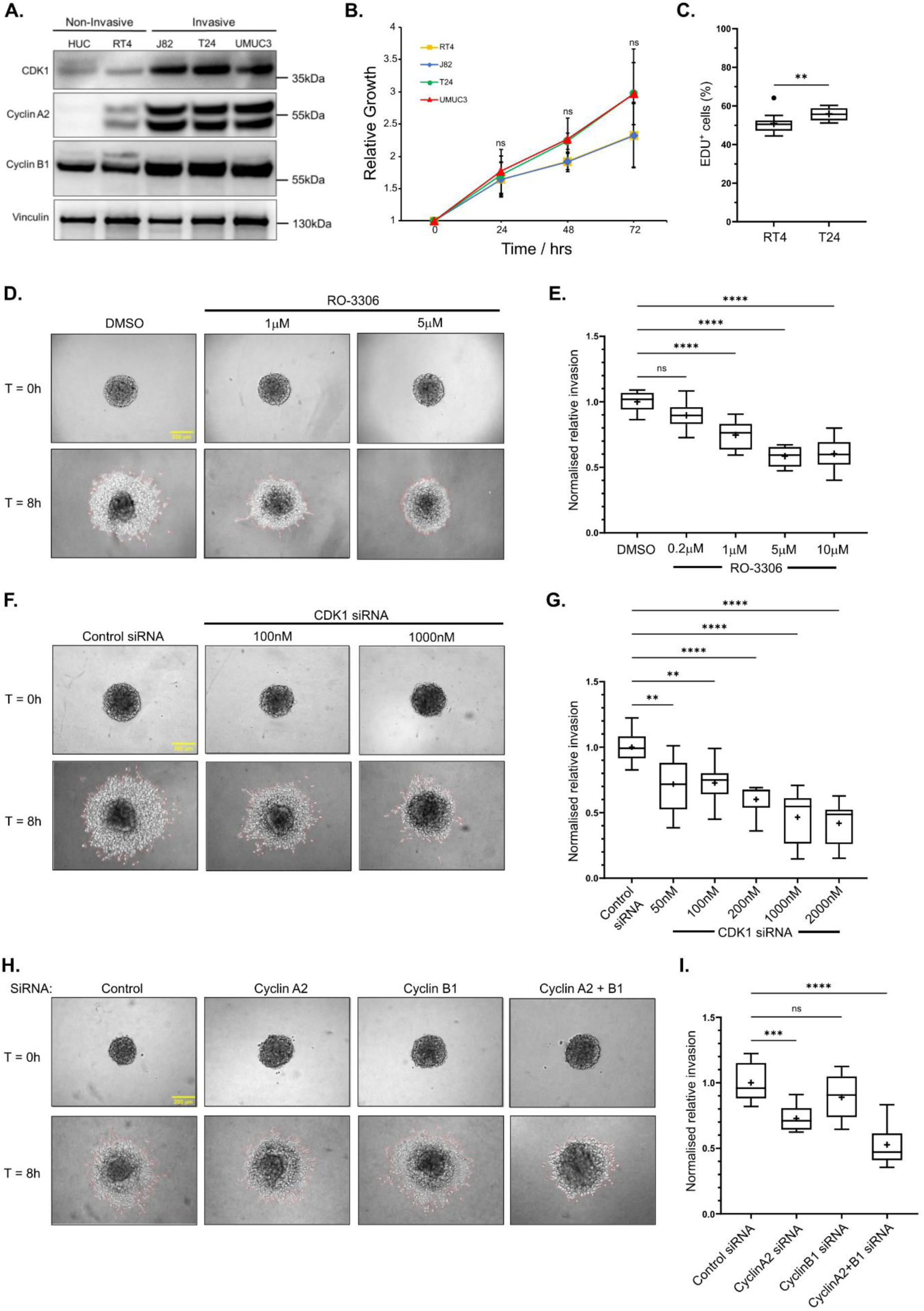
Loss of CDK1 and cyclin activity perturbs T24 bladder cancer cell invasion in 3D spheroid models. **A.** Western blot showing differences in expression of CDK1 and its respective cyclins cyclinA2 and cyclinB1, between non-invasive and invasive bladder cell lines. **B.** Quantification of growth curve analysis showing relative growth of bladder cell lines at 24 hour intervals for a total of 72 hours. Error bars show standard deviation. **C.** Quantification of EDU proliferation assay comparing the percentage of EDU positive cells between non-invasive RT4 and invasive T24 bladder cells. **D.** T24 spheroid invasion assay showing T24 spheroids seeded into 2.5mg/ml collagen gels supplemented with DMSO or varying concentrations of RO- 3306 at 0 hour and 8 hour timepoints (Only DMSO, 1uM RO-3306 and 5uM RO-3306 representative images shown). **E.** Quantification of invasion area of RO-3306 treated T24 spheroids relative to DMSO treated spheroids. A minimum of 9 spheroids per condition across 3 repeats was used for analysis. **F.** T24 spheroid invasion assay showing T24 spheroids generated from CDK1 knockdown T24 cells, seeded into 2.5mg/ml collagen gels at 0 hour and 8 hour timepoints (Only Control siRNA, 100nM CDK1 siRNA and 1000nM CDK1 siRNA representative images shown). **G.** Quantification of invasion area of CDK1 knockdown spheroids relative to control siRNA treated spheroids. A minimum of 9 spheroids per condition across 3 repeats was used for analysis. **H.** T24 spheroid invasion assay showing T24 spheroids generated from cyclinA2, cyclinB1 or a combination of cyclinA2+B1 knockdown T24 cells, seeded into 2.5mg/ml collagen gels at 0 hour and 8 hour timepoints. **I.** Quantification of invasion area of cyclin-knockdown spheroids relative to control siRNA treated spheroids. A minimum of 9 spheroids per condition across 3 repeats was used for analysis. One way ANOVA and Tukey’s post hoc test to compare between group means at 24, 48 and 72 hour timepoints used in B, unpaired student t-test used in C, one way ANOVA compared to control used in E, G and I, **** denotes p < 0.0001, *** denotes p < 0.001, ** denotes p < 0.01, ns denotes p > 0.05 (not significant).

### Cyclin-CDK1 control of migration and invasion is independent of the cell cycle

Having demonstrated that cyclin-CDK1 complexes are able to influence invasive cell migration using acute treatment with CDK1 inhibitors and longer-term depletion of CDK1 and cyclins, we next wished to determine whether the affects we were observing were due to a direct effect of CDK1 activity or an indirect effect of modulating the cell cycle. Treatment of cells with the CDK4/6 inhibitor Palbociclib results in a G0/G1 arrest that can be maintained over several days (Trotter and Hagan, 2020). Subsequently, this allows for knockdown of proteins to be undertaken in arrested cells, minimising any indirect effects of depleting CDK1, cyclinA2 and cyclinB1 may have on dividing cells. Short-term treatment of A2780 cells in CDMs or T24 cells in spheroid invasion assays (16 and 8 hours respectively) had no effect on cell motility or invasion (Figure 6A-D, Movie S3), demonstrating that CDK4/6 does not play a direct role in regulating cell migration in these contexts. Despite short-term treatment with Palbociclib altering the cell-cycle, complete arrest of cells in G1 did not occur (Figure S6) therefore we treated cells for 24 hours prior to undertaking CDK1 and cyclin knockdowns in the presence of Palbociclib. This longer-term treatment (80 hours) results in a complete G0/G1 arrest and in this context knockdown of CDK1 leads to a reduction in T24 cell invasion, demonstrating that CDK1 functions in G0/G1-arrested cells to regulate motility (Figure 6E-G). Interestingly, treatment with Palbociclib for 24 hours or above results in a reduction in cell invasion that is associated with a loss of CDK1, cyclinA2 and cyclinB1 (Figure 6E-G and Figure S6A). Taken together, these data suggest that the severe effect of CDK1- cyclin perturbation on cell motility does not extend to other cell cycle regulators such as CDK4/6, and that CDK1/cyclinA2/cyclinB1 have novel, non-canonical roles in migration and invasion which are independent of cell cycle phase. Furthermore, these data demonstrate that Palbociclib-induced G0/G1 arrest leads to a specific loss of cyclin-CDK1 complexes that potentially impacts upon the ability of cells to migrate in a 3D environment and, furthermore, highlights a role for CDK4/6 signalling in regulating CDK1 and cyclinA2/B1 expression levels.

**Figure 6:**
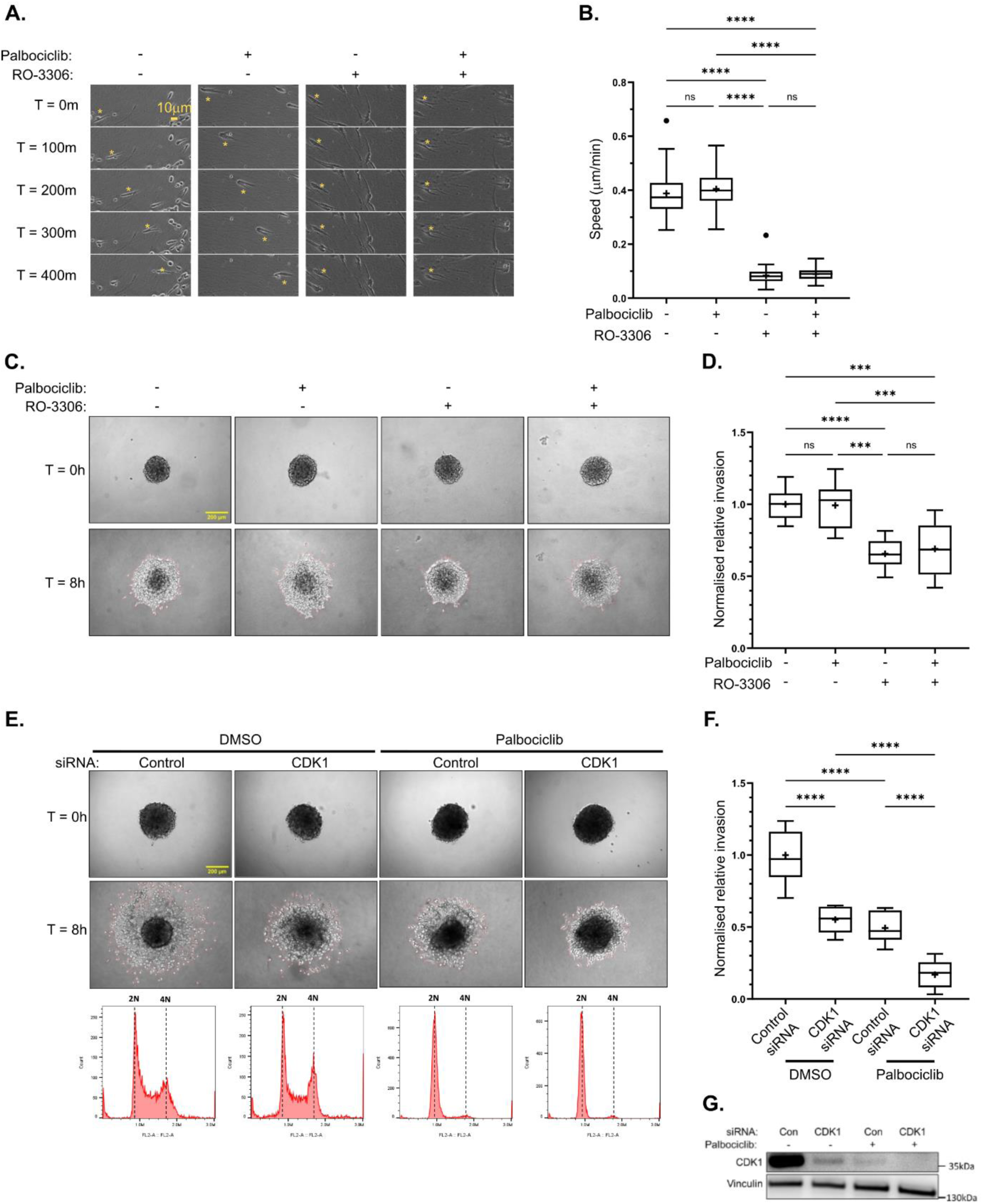
CDK1 regulates cell migration and invasion in G0/G1 arrested cells. **A.** A2780 cells seeded in CDM pre-treated (24 hours prior to imaging) with CDK4/6 inhibitor Palbociclib for arrest in G1 and/or CDK1 inhibitor RO-3306 (30 minutes prior to imaging) over 400 minute timelapse, yellow * denotes the position of the same cell at each time point. **B.** Average migration speed of A2780 cells in CDM treated with different CDK1 and CDK4/6 inhibitor combinations across 16 hour timelapse, n = 75 cells per condition analysed across 3 repeats. **C.** T24 spheroid invasion assay showing T24 spheroids seeded in 2.5mg/ml collagen gels pre-treated with Palbociclib and/or RO-3306 or DMSO for negative controls at 0 hour and 8 hour timepoints. **D.** Quantification of invasion of palbociclib and/or RO-3306 treated spheroids relative to DMSO treated controls. A minimum of 9 spheroids per condition across 3 repeats was used for analysis. **E.** T24 cells induced into G1 arrest by pre-treating with Palbociclib or DMSO (for controls) for 24h followed by control siRNA treatment or CDK1 knockdown, whilst maintaining cells in DMSO or Palbociclib (to maintain G1 arrest) for a total incubation time of 48 hours. Spheroids are generated from knockdown cells and incubated for 18-24 hours whilst maintaining G1 arrest using Palbociclib or DMSO for controls. Representative images show generated T24 spheroids seeded into 2.5mg/ml collagen gels pre-treated with DMSO or Palbociclib (to maintain G1 arrest over duration of experiment) at 0 hour and 8 hour timepoints. Histograms shows cell cycle profile of propidium iodide-stained cells used to generate spheroids for each condition. 2N and 4N highlight diploid and tetraploid cell populations, respectively. **F.** Quantification of invasion of CDK1 knockdown and palbociclib treated spheroids relative to control siRNA DMSO controls. A minimum of 9 spheroids per condition across 3 repeats was used for analysis. **G.** Western blot shows protein levels of CDK1 in control siRNA and CDK1 siRNA T24 cells pre-treated with DMSO or Palbociclib that were used for spheroid generation. One way ANOVA with multiple comparisons used in B, D and F, all comparisons shown in B and D, relevant comparisons of interest shown in F, **** denotes p < 0.0001, *** denotes p < 0.001, ns denotes p > 0.05 (not significant).

## Discussion

In summary, our major findings identify differential roles for cyclin-CDK1 complexes in regulating cell motility and invasion in 3D matrices in normal and cancer cells. CyclinB1-CDK1 plays a specific role in regulating Ect2 localisation and activation of RhoA at the rear of migrating cells, thereby facilitating the contractile feedback loop that drives rear cell movement during motility (Hetmanski et al., 2019). In contrast, cyclinA2-CDK1 plays a broader role in regulating cell motility via its localisation at both the protrusive front and the rear of migrating cells. Depletion of cyclinB1 or cyclinA2 results in differing cell morphologies with cyclinB1 depletion resulting in long, narrow cells indicative of a defect in rear retraction (Nguyen et al., 2018) whereas cyclinA2 depletion leads to the adoption of a large, rounded cell morphology. Knockdown of both leads to a combination of phenotypes that results in a complete abrogation of cell motility in addition to proliferation. These data define a role for cyclin-CDK1 complexes in regulating cell migration in addition to cell cycle progression. Furthermore, we observed increased expression of cyclinA2, cyclinB1 and CDK1 in invasive bladder cancer cells, in comparison to non-invasive cells, suggesting that modulation of cyclin-CDK1 complexes may play a significant role in facilitating invasive cell migration in cancer.

A role for cyclinA2 and CDK1 in cell migration has been described in a range of cell types, including breast and hepatocellular cancer cells and schwann cells (Chang et al., 2012; Fu et al., 2021; Lu et al., 2022) in addition to the cell types used in this study, demonstrating a conserved role for cyclinA2-CDK1 complexes in motility. In contrast, the role of cyclinB1-CDK1 in motility has not been previously described. We postulate that this is because cyclinB1-CDK1 plays a specific role in regulating the membrane tension-caveolae-RhoA contractility feedback loop at the rounded rear of cells that is only consistently observed in cells migrating within 3D matrices or in rigidity gradients (Hetmanski et al., 2019). The previous studies investigating the role of cyclins and CDK1 in migration have exclusively used 2D models of motility where the influence of cyclinA2-CDK1 on focal adhesions and the actin cytoskeleton dominates and the role of cyclinB1 may be minimal. This highlights the importance of considering how cell migration and invasion is regulated in 3D models.

We have demonstrated here that depletion of cyclinB1 or CDK1 perturbs localisation of the RhoA GEF Ect2 and subsequently RhoA activity at the rear of migrating cells and it will be interesting to determine whether direct phosphorylation of Ect2 by cyclinB1-CDK1 regulates Ect2 localisation and activity in motile cells. Ect2 is phosphorylated by cyclinB1-CDK1 during mitosis where it functions to activate RhoA to drive mitotic cell rounding and cytokinesis, as well as facilitating formation of the actin cortex in mitotic cells (Hara et al., 2006; Matthews et al., 2012; Niiya et al., 2006). We hypothesise that this role for cyclinB1-CDK1-dependent phosphorylation of Ect2 in regulating cortical actin and RhoA activity is conserved between mitotic and motile cells. Several other cytoskeletal regulators and focal adhesion proteins are phosphorylated during mitosis (Chen et al., 2022; Jones et al., 2019), suggesting that the conserved role of CDK1 in regulating the cytoskeleton is central to its function during mitosis, interphase and migration. Our data presented here and in our previous publications suggest distinct roles for cyclinB1- and cyclinA2- CDK1 complexes in cytoskeletal regulation, with the regulation of focal adhesions and phosphorylation of talin1 being primarily mediated by cyclinA2-CDK1 (Gough et al., 2021; Jones et al., 2018) and regulation of RhoA being driven by cyclinB1-CDK1. Identifying the cyclin-CDK1 complex specific mechanisms that regulate the cytoskeleton during cell cycle progression and migration within 3D environments therefore represents a significant avenue of future investigation.

In this study we demonstrate a role for cyclin-CDK1 complexes in regulating cell migration in normal RPE1 cells in addition to invasive migration in ovarian and bladder cancer cells, suggesting that modulation of cyclinA2, cyclinB1 and CDK1 expression or cyclin-CDK1 complex activity may contribute to the acquisition of invasive capabilities in cancer cells. Here, we show that CDK1, cyclinA2 and cyclinB1 are expressed at increased levels in invasive bladder cancer cells in comparison to non-invasive cells, therefore identifying the regulatory mechanisms that control CDK1 and cyclin expression is paramount. Overexpression of CDK1 is observed in several tumour types (Wang et al., 2023) yet the control of CDK1 expression levels is poorly understood. We have shown here that prolonged inhibition of CDK4/6 with palbociclib leads to a reduction in CDK1, cyclinB1 and cyclinA2. This suggests that CDK4/6 may play a role in regulating CDK1 and cyclin expression, or alternatively CDK1 and partner cyclins are specifically downregulated in quiescent cells. In this regard it is worth noting that a recent study using palbociclib to synchronise cells and study changes in protein expression during the cell cycle also observed a decrease in CDK1 in palbociclib treated cells, despite it being widely considered that CDK1 levels are consistent through the cell cycle (Rega et al., 2024). These observations also raise the possibility that the use of CDK1 and CDK4/6 inhibitors to treat invasive cancer types such as ovarian and bladder cancer might be a viable therapeutic option as they will simultaneously block both cell proliferation and invasive cell migration. Determining how cyclin-CDK1 complexes are regulated and function in cancer models such as patient-derived organoids and mouse models, and how therapeutically targeting CDK1 influences cancer progression and metastasis in these models represents an important area of future investigation.

## Methods

### Cell Culture and Transient Transfection

A2780 human ovarian cancer cells (female) and RT4, J82 and T24 human bladder cancer cells were maintained in RPMI-1640 medium (Sigma-Aldrich) containing L-Glutamine supplemented with 10% (v/v) fetal calf serum, and 1% (v/v) Antibiotic-antimycotic (Sigma-Aldrich); telomerase-immortalized fibroblasts (TIF) cells (used to produce CDMs) were maintained in Dulbecco’s modified Eagle’s medium (DMEM, Sigma-Aldrich) containing L-Glutamine and supplemented with 10% (v/v) fetal calf serum, and 1% (v/v) Antibiotic-antimycotic (Sigma Aldrich); human retinal pigment epithelial cells (RPE) cells were maintained in Dulbecco’s Modified Eagle Medium/Nutrient Mixture F-12 (DMEM/F12, Gibco) containing L-Glutamine and supplemented with 10% (v/v) fetal calf serum, 1% (v/v) 100x non-essential amino acid solution (Sigma Aldrich). Primary Human Urothelial cells (Sciencell) were maintained in Urothelial Cell Medium supplemented with urothelial cell growth supplement and 1% (v/v) penicillin-streptomycin. All cell lines were incubated at 37°C in a humidified 5% (v/v) CO2 atmosphere. siRNAs in A2780s and RPEs, and fluorescent constructs in A2780s, RPEs and T24s were transiently transfected by electroporation using a nucleofector (Amaxa, Lonza) using solution T, program A-23, 3μg DNA / 5μl 20 mM siRNA as per the manufacturer’s instructions. T24 cells were transfected with siRNAs using a Neon transfection system (Thermofisher) using Neon™ Resuspension Buffer R, 100μl Neon™ Tips, pulse voltage 1400 V, pulse width 20ms and pulse number 2, according to manufacturer’s instructions. Experiments were performed ∼72 hours after nucleofection for CCNB1 and CCNA2 and ∼48 hours after nucleofection for CDK1 for all cell types, and ∼48 hours after neon transfection for T24 cells.

### Reagents

Monoclonal antibodies used were mouse anti–cyclinA2 (clone BF683, 1:1,000 for WB; 4656; Cell Signaling Technology), mouse anti–CDK1 (clone POH1, 1:1,000 for WB; 4656; Cell Signaling Technology), rabbit anti-cyclinB1 (clone D5C10, 1:1,000 for WB, 1:100 for IF; 12231; Cell Signalling Technology); rabbit anti-cyclinD1 (clone E3P5S, 1:1,000 for WB; Cell Signalling Technology); rabbit anti-p27/KIP1 (clone D69C12, 1:1,000 for WB; Cell Signalling Technology) mouse anti-tubulin (clone DM1A, 1:10,000 for WB; ab7291; Abcam); mouse anti-vinculin (clone hVin-1, 1:2,000 for WB; V9264; Sigma-Aldrich). Secondary Alexa Fluor 680– conjugated (1:10,000; A10043; Thermo Fisher Scientific), DyLight 800–conjugated (1:10,000; 5257; Cell Signaling Technology) and HRP-conjugated (1:10,000; SA1-100 and SA1-200; Thermo Fisher Scientific) antibodies were used for immunoblotting. Anti–mouse and anti–rabbit Alexa Fluor 488–, 594–, and 647–conjugated secondary antibodies (1:1000) and Alexa fluor-tagged phalloidin were used for immunofluorescence (Thermo Fisher Scientific). Palbociclib and RO-3306 were purchased from Sigma-Aldrich, Flipper-TR was purchased from Spirochrome. The following plasmids used in this study were obtained as gifts: FRET biosensor Raichu-1237X RhoA (Yoshizaki et al., 2003) was kindly provided by Prof. M. Matsuda; mCherry-Caveolin-1 construct (Hayer et al., 2010) was kindly provided by Dr M. Bass; Emerald-Lifeact was kindly provided by Prof. C. Ballestrem; BFP-Ezrin constitutively active (CA) (Gautreau et al., 2000) was kindly provided by Prof. E. Paluch; GFP-Cavin-1 was kindly provided by Dr J. Goetz; Lifeact-7-iRFP670 was a gift from Ghassan Mouneimne (Addgene plasmid #103032); pLenti Lifeact-mTagBFP2 PuroR was a gift from Ghassan Mouneimne (Addgene plasmid # 101893). SiRNAs used were: CDK1, SMARTpool reagent L-003224-00- 0005, Horizon Discovery; cyclinB1 SMARTpool reagent L-003206-00-0005, Horizon Discovery; CCNA2, validated silencer select oligonucleotide s2514, Thermo Fisher Scientific; CCNB2 validated silencer select oligonucleotide s2517, Thermo Fisher Scientific.

### CDM generation

Cell derived matrices were generated using the method developed by the Yamada lab (Caswell et al., 2007; Cukierman et al., 2001). 6/12-well plastic (Corning) or 35 mm glass bottom (Mattek) plates were prepared by coating with 0.2% gelatin (v/v, Sigma Aldrich) for 1 hour, crosslinking with 1% glutaraldehyde (v/v, Sigma Aldrich) for 30 minutes and quenching with 1M glycine (Thermo Fisher) for 20 minutes before TIFs were confluently seeded. DMEM medium supplemented with 0.25% ascorbic acid (v/v, Sigma Aldrich) was changed every 48 hours for 8 days. Cells were denuded with extraction buffer (20 mM ammonium hydroxide (NH_4_OH); 0.5% (v/v) Triton X-100) to leave only matrix before treatment with 10μg/ml DNAse 1 (Roche) to cleave phosphodiester linkages in the DNA backbone. CDMs were stored at 4°C with 1% (v/v) anti-biotic anti-mycotic (Sigma) and used within 3 months of generation.

### CDM migration

A2780/RPE cells were seeded at sparse (∼50,000 cells/well) confluency in 6-well CDMs and allowed to spread for ∼ 4 hours. Images were acquired on an Eclipse Ti inverted microscope (Nikon) using a 20x/ 0.45 SPlan Fluar objective, the Nikon filter sets for Brightfield and a pE-300 LED (CoolLED) fluorescent light source with imaging software NIS Elements AR.46.00.0. Point visiting was used to allow multiple positions to be imaged within the same time-course and cells were maintained at 37°C and 5% CO2. The images were collected using a Retiga R6 (Q-Imaging) camera. 5 randomly chosen positions per cell were captured every 10 minutes over 16 hours (or 60 hours for A2780 cyclinA2+B1 siRNA experiment). 5 randomly chosen cells per position (meaning 25 cells tracked per condition per experiment) were individually manually tracked over the entire 16 hours or first 16 hours of the timelapse using the ImageJ plugin MTrackJ every 3 frames (i.e. using 30 min timepoint intervals). Representative images of individual cells/fields of view are shown where appropriate.

### Live fluorescent imaging

All live fluorescent images were acquired using a CSU-X1 spinning disc confocal (Yokagowa) on a Zeiss Axio-Observer Z1 microscope with a 63x/1.40 Plan-Apochromat objective for CDMs and a 40x/1.40 Plan-Apochromat objective for collagen gels. An Evolve EMCCD camera (Photometrics) and motorized XYZ stage (ASI) was used. The 405, 488, 561 and 647nm lasers were controlled using an AOTF through the laserstack (Intelligent Imaging Innovations (3I)) allowing both rapid ‘shuttering’ of the laser and attenuation of the laser power. Images were captured using SlideBook 6.0 software (3i). Randomly chosen representative polarized Lifeact-Emerald + mCherry-Caveolin-1, or Lifeact-iRFP670 expressing cells were captured with the appropriate excitation/emission spectrum and exposure time following ∼ 4 hour spreading time in CDM and ∼ 4 hour spreading time in collagen gels in 1x Opti-Klear medium (Marker Gene Technologies Inc) supplemented with 10% (v/v) FCS. Where indicated, cells were dyed with Hoechst 33258 for 10 minutes prior to imaging. Cells were imaged every 30s for 5 minutes, and rear translocation in CDM was measured for all imaged cells in ImageJ across this 5 minute period by comparing the position of a part of the rear of the cell at the start of the timelapse with the exact same part of the rear at the end of the timelapse. Rear to nucleus distance was determined in ImageJ by measuring the distance between the rearmost point of the nucleus to the rearmost point of the membrane at the first time interval. For cell size/shape metrics in collagen gels, the ‘width’ and ‘length’ of the cells were manually determined at the widest and longest single positions in the cell, while aspect ratio is defined as length/width. For nucleus size, the Hoechst 405 channel was thresholded in ImageJ to create the nuclear mask then the size of the mask determined by the measurement command. All cells in collagen gels were imaged live to determine polarity and movement direction and analysis was done on the 1_st_ frame captured.

### Staining/fixed imaging

Cells were fixed in 4% paraformaldehyde (PFA) at room temperature following ∼ 4h spreading in CDM after non-confluent (∼50,000 cells/plate) seeding. For osmotic shock, medium was replaced with 50% 1x RPMI, 50% distilled water 30 minutes prior to fixation, while medium was replaced at the same time with 100% 1x RPMI in the control isotonic condition. Membranes were permeabilized with 0.2% (v/v) Triton-X and blocked in 5% (w/v) heat-denatured bovine serum albumin (BSA) before being stained with appropriate antibodies as in reagents section. Cells were incubated with secondary antibodies Rabbit Alexa Fluor 647-conjugated and Mouse Alexa Fluor 594 conjugated for 1 hour, (both Invitrogen), stained with Phalloidin Alexa Fluor 488-conjugated (Invitrogen) for 1 hour, and Hoechst 33258 (ThermoFisher) for 10 minutes.

Cells were imaged using a Leica TCS SP8 STED microscope with an HC PL APO 100 x/1.40 oil objective using a HyD1 detector, while a notch filter was used for background reduction where possible. Images were captured using a white light laser (WLL) with excitation wavelengths 488, 561, 633 nm and appropriate emission spectra for green, red and far red respectively, while a Diode 405 laser was used with appropriate emission spectra for blue. Z-stacks were captured covering the entire Z-profile of the cell with intervals of 0.3 μm at a zoom of 1x. All subsequent analysis and quantification was performed on maximum intensity projections (MIPs) in ImageJ; all representative images shown throughout are MIPs (pseudocolored using the red hot LUT where appropriate to reveal differences in intensity).

For quantification of localized CDK1 and cyclinB1 intensity, images were analysed with the interactive 3-D surface plot v2.31 in ImageJ using a 128 grid size and smoothing factor of 3.0. Under these conditions, the peak intensities of the rear, front and nuclear cell regions were manually identified, and the ratio rear peak / non-rear peak or rear peak / nuclear peak calculated, therefore providing inherent normalization avoiding staining/expression discrepancies. The rear peak / non-rear peak ratios between conditions were then directly compared.

### Collagen gel generation for single cell imaging

Collagen gels were generated using an adapted approach to collagen gel generation used in inverted invasion assays described previously (Hennigan et al., 1994; Hetmanski et al., 2016). 10x RPMI, water, NaHCO_3_, NaOH were mixed and added in equal volume to 3mg/ml collagen (Gibco) and supplemented with fibronectin, while being kept on ice to prevent polymerisation. Polymerised rat tail collagen was labelled with Alexa-488 NHS Ester before being solubilised and added at a 1:10 ratio with unlabelled collagen. ∼50,000 cells of interest per condition (A2780/RPEs previously transfected with siRNA as above) were centrifuged at 1400 RPM for 4 minutes and seeded directly into the collagen gel mix (while still liquid, pre polymerisation). 50μl of the cells/collagen mixture was pipetted onto the glass area of a 35mm Mattek dish (to cover the circular glass surface) and the collagen was allowed to polymerise for 30 minutes at 37°C. Dishes were then gently agitated to detach the gels from the glass area (to ensure cells in 3D collagen and not adhered to the glass surface below) and normal growth medium added. Cells were imaged 16 hours post seeding. For Flipper-TR experiments, Flipper-TR was added at 1:250 following collagen addition (prior to cell seeding/polymerisation) and cells were imaged ∼4 hours post seeding.

### RhoA FLIM imaging

Cells previously transfected with GFP-RFP Raichu RhoA were seeded into CDM 4h prior to imaging and imaged at 37°C in 1x OptiKlear with 10% FCS. Cells were imaged on a Leica SP8 gSTED microscope using Leica FALCON hardware/software for fluorescent lifetime imaging (FLIM) of the GFP donor channel with 488 nm excitation and 498-550 emission with speed 50Hz, line average 6, and format 512x512 pixels. Following imaging, images were subjected to 2x2 binning (to improve signal to noise). For quantification, regions at the cell rear and cell front were manually identified and drawn in the Leica FALCON software, and the mean average lifetimes measured by fitting a single exponential to these front/rear ROIs. For the lifetime representative images, the lifetime limits were set to 1-2ns, the counts limits to 0-2000 and the images exported with the default Leica LUT.

### FlipperTR FLIM imaging

Cells were seeded directly into collagen gels containing 4μM Flipper TR 4 hours prior to imaging and imaged at 37°C in 1x OptiKlear (without FCS). Cells were imaged on a Leica SP8 gSTED microscope using Leica FALCON hardware/software for fluorescent lifetime imaging (FLIM) with 488 nm excitation and 575-625 emission with speed 50Hz, line average 6, and format 512x512 pixels. Following imaging, images were subjected to 2x2 binning (to improve signal to noise) and a double exponential was fit to ensure a close fit (chi squared < 1.5). For quantification, membrane regions at the cell rear and cell front were manually identified and drawn in the Leica FALCON software, and the mean average lifetimes measured by fitting a single exponential to these front/rear ROIs. For the lifetime representative images, the lifetime limits were set to 3-5ns, the counts limits to 0-1000 and the images exported with the default Leica LUT.

### Inverted invasion assay

Inverted invasion assays were performed based on the protocol as described previously [Hennigan et al., 1994]. ‘Stiff’ (final concentration 5 mg/ml; Corning) or ‘soft’ (final concentration 1.5mg/ml, Gibco) Collagen I supplemented with 25 μg/ml fibronectin was allowed to polymerise in inserts (Transwell; Corning) for 1 hour at 37°C upon addition mixing with 10x RPMI, water, NaHCO3 and NaOH to regulate pH. Transwells were then inverted and ∼50,000 cells were seeded directly onto the bottom surface for 4 hours at 37°C. Transwells were then re-inverted, washed and placed in serum free medium. Medium supplemented with 10% FCS and 30 ng/ml EGF, was placed on top of the matrix to provide a chemotactic gradient for invasion. After 72 hours, all cells were stained with Calcein-AM ∼1 hour prior to imaging and visualised by confocal microscopy with serial optical sections being captured at 15-μm intervals using an inverted confocal microscope (SP8 gSTED, Leica) using a 20× objective. Invasion was quantified using the area calculator plugin in ImageJ, where the invasive proportion was obtained by measuring the fluorescence intensity of cells invading > 45 μm the 5th Z-stack onwards and dividing this by the total fluorescence intensity in all Z-stack images (15 planes in total).

### Immunoblotting

Cells were lysed in lysis buffer (200nM NaCl; 75mM Tris; 15mM NaF; 1.5mN Na_3_VO_4_; 7.5mM EDTA; 7.5mM EGTA; 1.5% (v/v) Triton X-100; Igepal CA-630) supplemented with Halt_TM_ protease (100x) and phosphatase (100x) inhibitor cocktail (Thermo Fisher). Lysates were clarified by centrifugation at 10,000 g for 20 minutes at 4°C.

Cell lysates were separated by SDS-PAGE (4-12% Bis-Tris gels; Thermo Fisher Scientific) under reducing conditions and transferred onto nitrocellulose membranes (Cytiva). Membranes were blocked for 60 minutes at RT using casein-blocking buffer (Sigma-Aldrich) and then probed overnight with primary antibodies diluted in TBST (110mM Tris-HCl, pH 7.4, 150mM NaCl, 0.05% Tween-20) at 4°C. Membranes were washed for 15 minutes by using TBST and then incubated with the appropriate secondary antibodies diluted in TBST for 90 minutes at RT. Membranes were washed for 15 minutes by using TBST and exposed to chemiluminescent substrate and bound antibodies were visualised using a Syngene G:BOX CHEMI XRQ (Syngene) for T24, RT4, HUC, J82 and UMUC3 lysates; or scanned using an infrared imaging system (Odyssey; LI-COR Biosciences) for A2780 and RPE lysates. Band intensities normalised to loading controls were analysed using ImageJ throughout.

### 3D spheroid invasion assay

Spheroids were generated using RT4 and T24 bladder cells utilizing the hanging-drop method. Cells were suspended in complete RPMI and methocel solution (12mg/ml methylcellulose diluted in complete RPMI) at a 4:1 ratio. Cell suspensions were added to the under-side of a petri dish lid in 20 μl droplets at a density of 2000 cells per droplet and inverted onto the base of a petri-dish containing 5ml PBS to prevent spheroids from drying out. Cells aggregated on the bottom of 20 μl droplets due to gravity and formed compact spheroids after 18-24 hours. Collagen gels were generated using PureCol™ EZ Gel (Sigma Aldrich) by diluting in neat RPMI to a concentration of 2.5mg/ml and stored at 4°C. In a 96-well plate, 40 μl of 2.5mg/ml collagen gel was added to each well and incubated at 37°C for 24 hours alongside spheroids to allow for sufficient gelation to ensure spheroids wouldn’t sink and form a monolayer on the bottom of the well. Following spheroid generation, an additional 60 μl of 2.5mg/ml collagen gel was added to initial 40 μl gel layers and spheroids in hanging droplets were removed from droplets in a volume of 5 μl using a p20 pipette and injected directly into 60 μl collagen gel layer. Spheroids in gels were first imaged at 20x magnification using a DMi8 inverted microscope (Leica) and then incubated for 90 minutes to allow sufficient gelation, following which 100 μl of complete RPMI was added to each gel. Spheroids were then incubated further at 37°C to a total incubation time of 8 hours before being imaged again using the DMi8 inverted microscope. Area of invasion was measured using Image J and determined by measuring initial spheroid area at 0 hour and subtracting value from total area following 8 hour invasion into collagen gel.

For CDK drug treatments, collagen gels were additionally supplemented with either DMSO, 2 μM Palbociclib or 10 μM RO-3306 (1:1000 dilution from stock solutions).

### Growth curve analysis

Bladder cells were seeded into wells of a 96-well plate at a density of 2.5x10^3^ and incubated at 37°C for up to 72 hours. At indicated time-points (0, 24, 48 and 72 hours), cells were pulsed with 4 μM Calcein-AM (Dojindo) and incubated at 37°C for 1 hour to allow for sufficient esterase activity. The number of viable cells was determined by measuring fluorescence at an excitation wavelength of 485nm and an emission wavelength of 520nm using a FLUOstar Omega Microplate reader (BMG Labtech).

### Flow cytometry

For cell cycle analysis, cells were washed once with phosphate-buffered saline (PBS) and released from the substrate with trypsin. Cells were pelleted and washed in PBS before fixation in ice-cold Ethanol and stored at -20°C for a minimum of 24 hours. Fixed cells were pelleted and washed three times in PBS at RT before final pellet was resuspended in 500 μl FxCycle™ PI/RNase Staining Solution (Thermo Fisher) and incubated at RT for a minimum of 30 minutes. Samples were ran on a Accuri™ C6 Flow Cytometer (BD Biosciences), with a total of 1x10^4^ cells counted for each sample. Data were analysed with FlowJo™ software (BD Biosciences).

### EDU Proliferation Assay

Bladder cells were cultured on coverslips at a density of 4.5 x10^4^cells and incubated at 37°C for 24 hour. Following 24 hour incubation, cells were pulsed with 10 μM EDU and incubated at 37°C for a further 2 hours. The prepared cells were washed three times in PBS and fixed with 4% paraformaldehyde for 15 minutes at room temperature, followed by membrane permeabilization in 0.2% Triton X-100/PBS for 10 min and blocking with 1% BSA supplemented with 0.1M Glycine to quench fixative for 30 minutes. Cells were EdU-labelled using Click-it chemistry according to the manufacturer’s instructions (Thermo Fisher) and counter stained with Hoechst 33342 (Thermo Fisher). Images were acquired using a Leica SP8 confocal microscope with a 20x objective and Leica LAS X Software (Leica). Image analysis was performed using ImageJ.

## Supplementary Movies

Movie S1: Timelapse movie of control (left), cyclinA2 (centre) and cyclinB1 (right) siRNA A2780 cells moving in CDM over 16 hour timelapse.

Movie S2: Timelapse movie of control (left), and cyclinA2+B1 (right) siRNA A2780 cells moving in CDM over 60 hour timelapse.

Movie S3: Timelapse movie of cells moving in CDM over 16 hour timelapse treated with RO- 3306 and Palbociclib as indicated (DMSO used for control)

**Figure S1:**
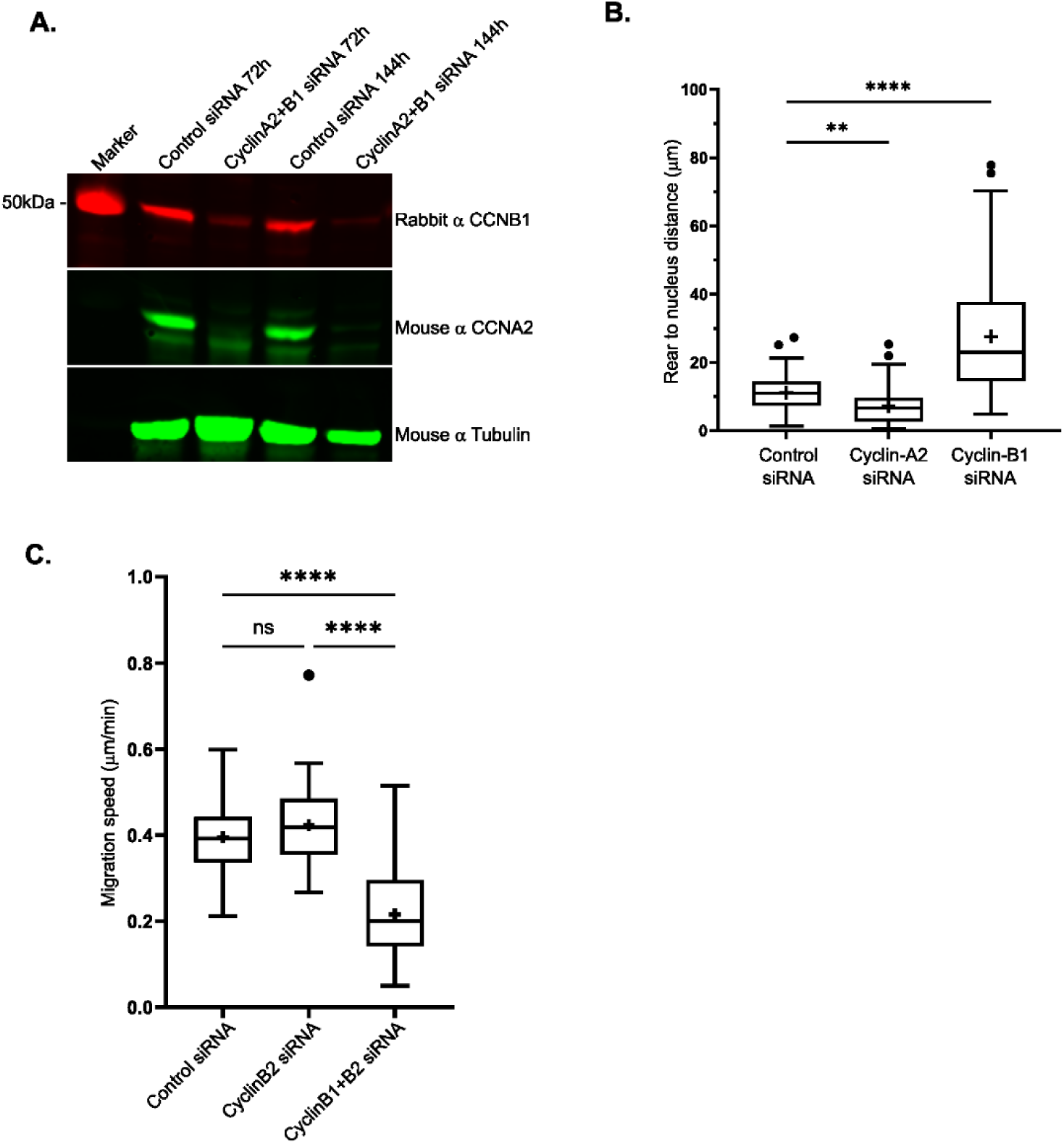
Knockdown of cyclinA2 and cyclinB1 alters nuclear positioning in migrating cells and knockdown of cyclinB2 has no effect on cell migration. **A.** Western blot showing cyclinA2 (CCNA2), cyclinB1 (CCNB1) and Tubulin (for loading) protein levels for control siRNA and cyclinA2+B1 siRNA A2780 samples corresponding to the start of the longterm timelapse imaging (72h) and at the end of the longterm timelapse imaging (144h). **B.** Average distance between the nucleus and cell rear of control, cyclinA2 and cyclinB1 siRNA A2780 cells (as determined using the lifeact channel overexposed and the rearmost nucleus and rearmost membrane points in the first frame), n > 40 cells per condition across 3 repeats. **C.** Average migration speed of cyclinB2 and concomitant cyclinB1+B2 siRNA A2780 cells in CDM across 16 hour timelapse, n = 50 cells per condition analysed across 3 repeats. One way ANOVA compared to control used in B and C, **** denotes p < 0.0001, ** denotes p < 0.01, ns denotes p > 0.05 (not significant).

**Figure S2:**
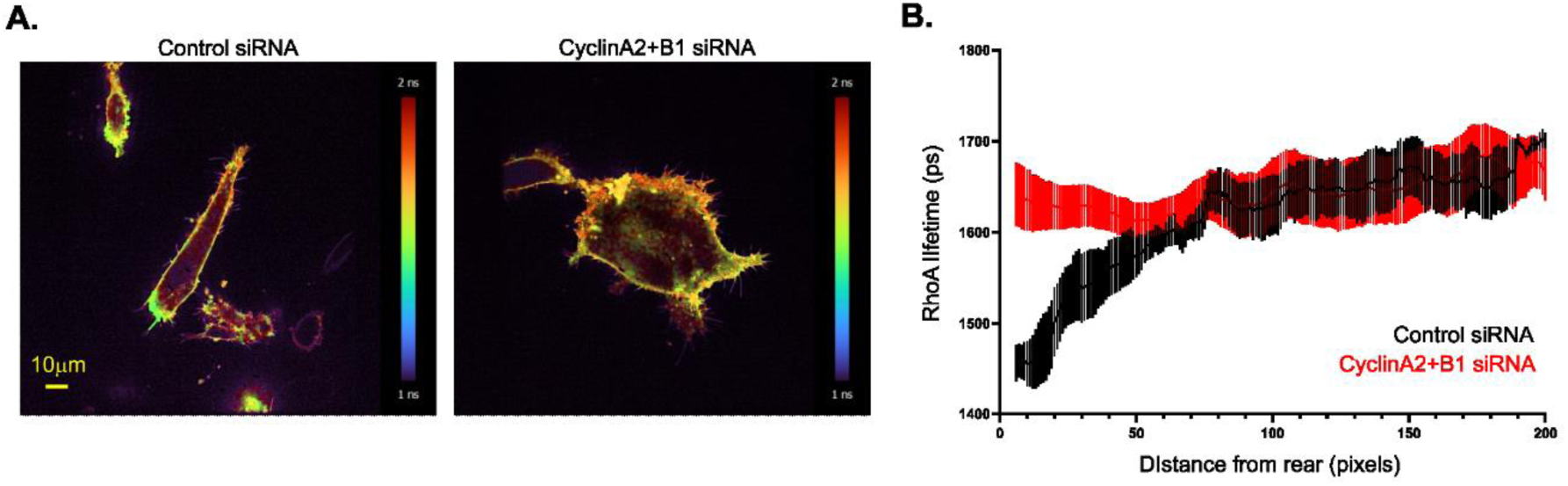
Knockdown of cyclinA2 and cyclinB1 perturbs activation of RhoA at the rear of migrating cells. **A.** Control and cyclinA2 + cyclinB1 concomitant knockdown A2780 cells seeded in CDM transfected with GFP-RFP Raichu-RhoA probe, FLIM lifetime of donor GFP channel shown, blue denotes shorter lifetime (high activity), yellow/red denotes higher lifetime (low activity), colourbar shown. **B.** Average lifetime across the entire width of the membrane per pixel from the rear, (averaged to nearest 10 front-rear length pixels) for control (black line/bars) and cyclinA2 + cyclinB1 siRNA (red line/bars) A2780s in CDM; standard error mean (SEM) shown, averaged across >12 cells for each condition.

**Figure S3:**
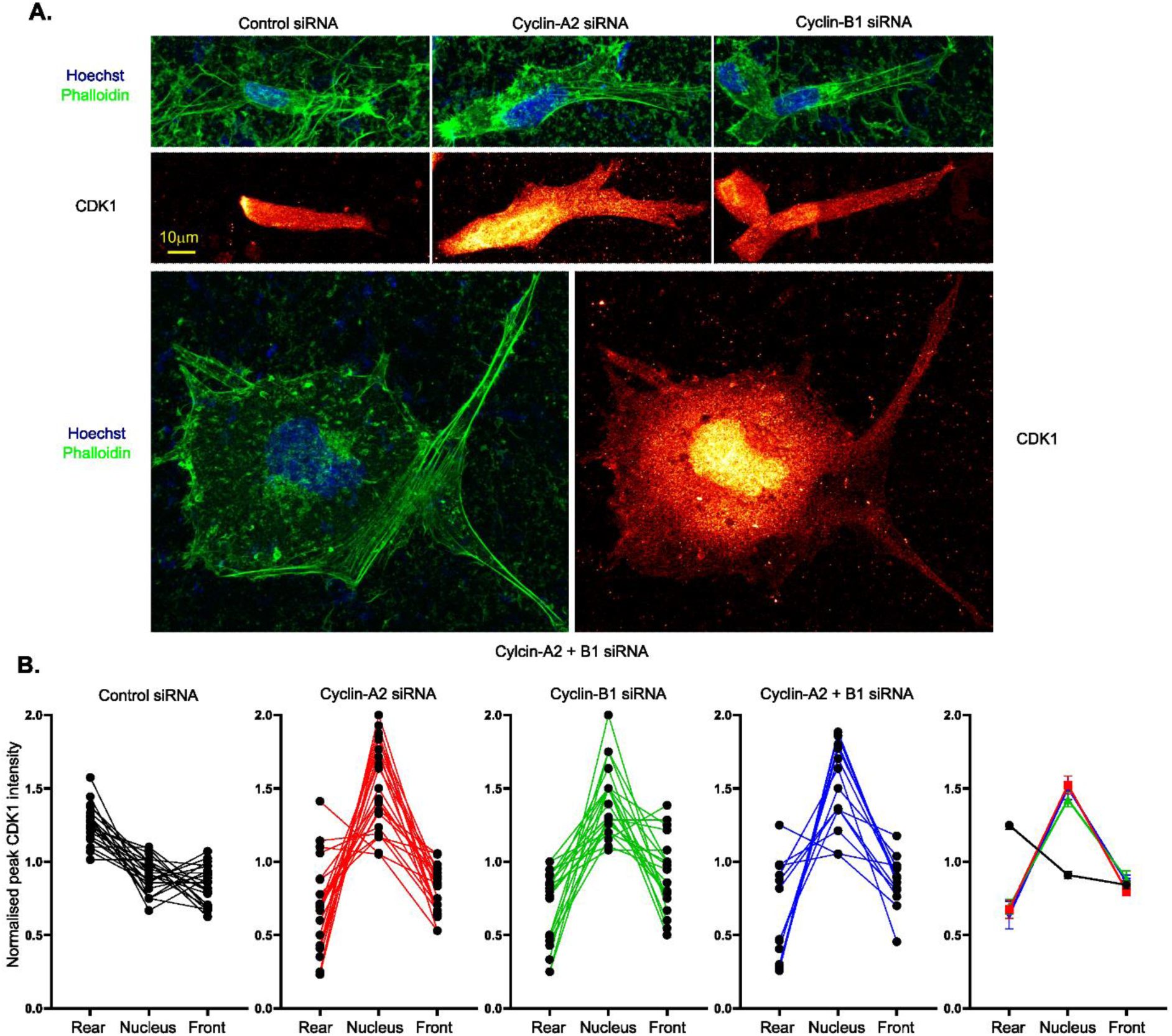
Knockdown of cyclinA2 or cyclinB1 perturbs CDK1 rear localisation in migrating cells. **A.** Control, CyclinA2, CyclinB1, and cyclinA2 + cyclinB1 concomitant knockdown A2780 cells fixed 4 hours post seeding in CDM, stained with 488-phalloidin for actin (green, top/left images), Hoechst 33258 for nucleus (blue, top/left images) and CDK1 (Mouse α CDK1 primary, α Mouse Cy3 secondary, red hot look- up table applied, bottom/right images). **B.** Normalised CDK1 staining intensity in rear, nucleus and leading edge manually identified regions of A2780s in CDM; left 4 graphs for each condition as indicated where each connected line represents a single cell, right graph shows collated data from all cells for each condition with error bars representing SEM.

**Figure S4:**
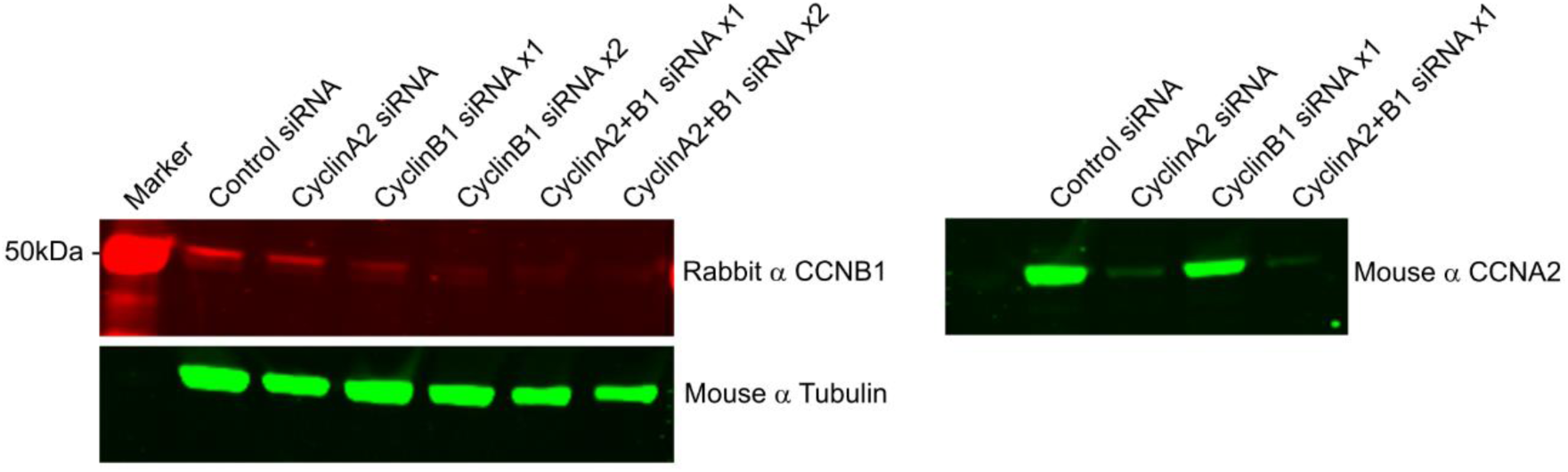
Representative Western blots showing cyclin knockdowns in RPE cells. Top: Labelled western blot showing cyclinA2 (CCNA2), cyclinB1 (CCNB1) and Tubulin (for loading) protein levels for control, cyclinA2, cyclinB1 and cyclinA2+B1 siRNA RPE samples corresponding to the start of the longterm timelapse imaging (72h), x1 denotes single knockdown, x2 denotes knockdown performed on consecutive days 24 hours apart.

**Figure S5:**
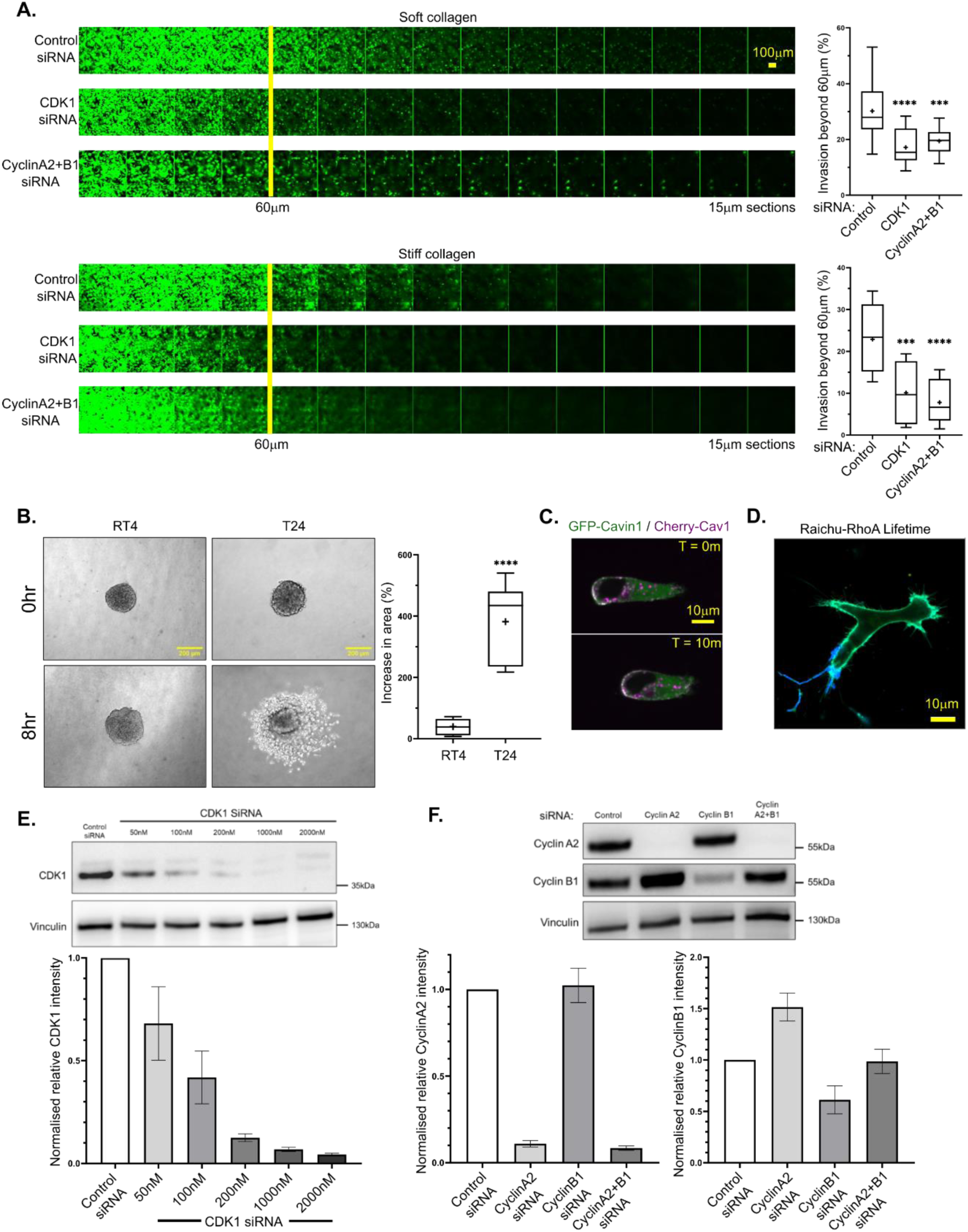
Knockdown of CDK1, cyclinA2 and cyclinB1 perturb ovarian and bladder cancer cell invasion in 3D collagen gels. **A.** Left: Representative montage images of Z planes from inverted invasion assay using A2780 cells stained with calcein-AM 1h prior to imaging, each image- taken 15mm apart whereby cells which are visible beyond the 4_th_ slice (60mm) are taken as invasive for cells in ‘soft’ (final concentration 1.5mg/ml, top) and ‘stiff’ (final concentration 5mg/ml, bottom) collagen; right: Quantification of % of cells which have invaded beyond 60mm, calculated as the total thresholded intensity in beyond the 4_th_ slice / the total thresholded intensity of the whole stack, n=9 fields analysed per condition across 3 repeats. **B.** Spheroids generated from invasive T24 or non-invasive RT4 cells seeded into 2.5mg/ml collagen gels at 0h and 8h timepoints. Quantification (right) of percentage increase in area of T24 spheroids relative to RT4 spheroids following 8h incubation, n=9 spheroids analysed per condition across 3 repeats. **C.** Representative T24 cell transfected with GFP-Cavin1 (green) and cherry-caveolin-1 (magenta, white overlap indicated caveolae localisation) seeded in CDM at two time points 10 minutes apart. **D.** Representative T24 cell transfected with GFP-RFP Riachu-RhoA, lifetime of FLIM image of GFP donor channel shown where blue denotes shorter lifetime, higher RhoA activity. **E.** Western blot showing changes in CDK1 protein levels of T24 cells following treatment with varying concentrations of CDK1 siRNA used in spheroid invasion assay. Quantification (below) of CDK1 protein levels relative to control siRNA normalised to vinculin. Error bars show SEM. **F.** Western blot showing cyclinA2 and cyclinB1 protein levels in T24 cells following treatment with control, cyclinA2, cyclinB1 or a combination of cyclinA2+B1 siRNA. Quantification (below) of cyclinA2 (bottom/left) and cyclinB1 (bottom/right) protein levels relative to control siRNA normalised to vinculin. Error bars show SEM.

**Figure S6:**
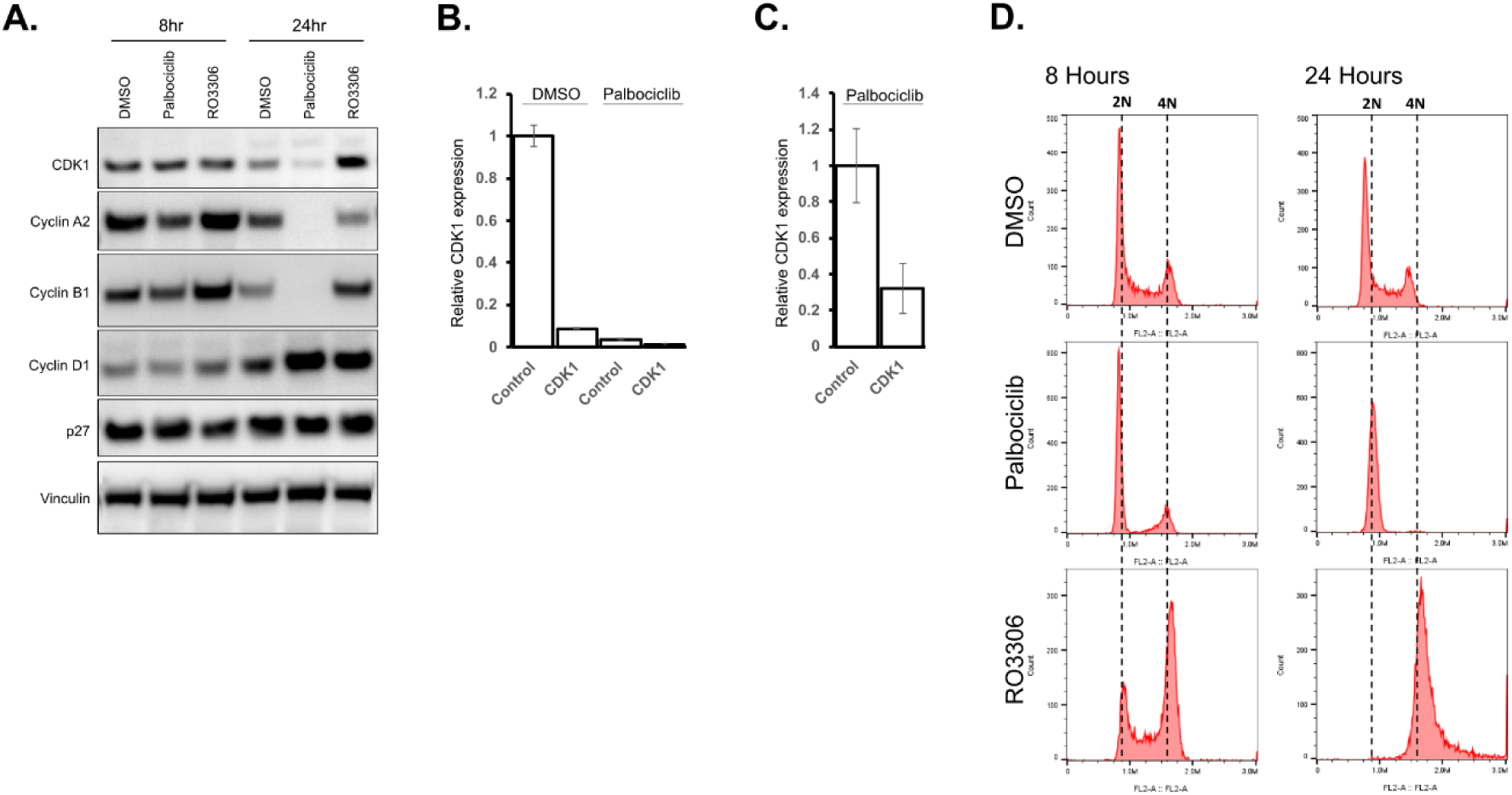
Treatment with palbociclib for 24 hours leads to a reduction in CDK1, cyclinA2 and cyclinB1. Knockdown of CDK1 and cyclins leads to a further reduction in invasion in palbociclib treated cells. **A.** Western blot showing CDK1, cyclinA2, cyclinB1, cyclinD1, p27 and Vinculin (for loading) protein levels for T24 cells following 8h and 24h treatment with DMSO, Palbociclib or RO-3306. **B.** Quantification of western blot from Figure 6C. highlighting CDK1 protein levels of CDK1 knockdown T24 cells treated with DMSO or Palbociclib relative to DMSO-treated control siRNA cells. Error bars show SEM **C.** Quantification of western blot from Figure 6C. highlighting relative CDK1 protein levels of palbociclib treated cells only. Error bars show SEM **D.** Flow cytometry analysis showing cell cycle profiles of T24 cells following 8h and 24h treatment with DMSO, Palbociclib or RO-3306 using propidium iodide-stained cells.

